# Non-canonical proline-tyrosine interactions with multiple host proteins regulate Ebola virus infection

**DOI:** 10.1101/2020.05.19.102954

**Authors:** Jyoti Batra, Manu Anantpadma, Gabriel I. Small, Olena Shtanko, Mengru Zhang, Dandan Liu, Caroline G. Williams, Nadine Biedenkopf, Stephan Becker, Michael L. Gross, Daisy W. Leung, Robert A. Davey, Gaya K. Amarasinghe, Nevan J. Krogan, Christopher F. Basler

**Affiliations:** J. David Gladstone Institutes, San Francisco, CA 94158, USA; Department of Microbiology, NEIDL, Boston University School of Medicine, Boston, MA, 02118, USA; Department of Pathology and Immunology, Washington University School of Medicine, St. Louis, MO 63105, USA; Host-Pathogen Interactions, Texas Biomedical Research Institute, San Antonio, TX 78245, USA; Department of Chemistry, Washington University School of Medicine, St. Louis, MO 63105, USA; Center for Microbial Pathogenesis, Institute for Biomedical Sciences, Georgia State University, Atlanta, GA 30303, USA; Institute of Virology, Philipps University of Marburg, Marburg, Germany; Department of Cellular and Molecular Pharmacology, University of California, San Francisco, CA 94158, USA; Quantitative Biosciences Institute, University of California, San Francisco, CA 94158, USA; John T. Milliken Department of Medicine, Division of Infectious Diseases, Washington University School of Medicine, St. Louis, MO 63110, USA

**Keywords:** Ebola virus, virus-host interactions, VP30, RNA viruses, viral replication

## Abstract

The Ebola virus VP30 protein interacts with the viral nucleoprotein and with host protein RBBP6 via PPxPxY motifs. In these interactions the largely alpha-helical carboxy-terminal domain of the EBOV VP30 engages with the motif such that the prolines adopt non-canonical orientations, as compared to other proline-rich motifs. Affinity tag-purification mass spectrometry identified additional PPxPxY-containing host proteins, including hnRNP L, hnRNPUL1 and PEG10, as VP30 interactors. Of these, hnRNP L and PEG10, like RBBP6, inhibit viral RNA synthesis and EBOV replication, whereas hnRNPUL1 enhances. Further, double knockdown studies support additive effects of RBBP6 and hnRNP L. Binding studies demonstrate variable capacity of PPxPxY motifs to bind VP30 and the extended motif PxPPPPxY is demonstrated to confer optimal binding and to inhibit RNA synthesis, with the fifth proline and the tyrosine being most critical. Competition binding and hydrogen-deuterium exchange studies demonstrate that each protein binds a similar interface on VP30 and impacts VP30 phosphorylation. VP30 therefore represents a novel proline recognition domain that allows multiple host proteins to target a single viral protein-protein interface to modulate viral transcription.

## Introduction

*Zaire ebolavirus* (Ebola virus or EBOV), a member of the filovirus family of enveloped, non-segmented, negative-sense RNA viruses, is a zoonotic pathogen notable for its propensity to cause outbreaks of severe disease in humans. The public health significance of EBOV is evidenced by past outbreaks where reported case fatality rates ranged up to 90 percent; by the West Africa EBOV epidemic in 2013-2016, which resulted in more than 28,000 infections and more than 11,000 deaths and export of infected individuals from West Africa to other areas of the world; and by three outbreaks in the Democratic Republic of Congo that have occurred from 2017-2020 (Bausch & Rojek, 2016, Ilunga Kalenga, Moeti et al., 2019, Nsio, Kapetshi et al., 2019).

The filovirus genomic RNA is ~19 kilobases in length and has seven separate transcriptional units (genes) which encode distinct mRNAs. The genome is encapsidated by nucleoprotein (NP) with the resulting ribonucleoprotein serving as the template for RNA synthesis reactions that replicate the viral genomic RNA and transcribe the mRNAs that lead to viral gene expression. Viral genome replication requires, in addition to NP, the viral proteins VP35 and L, the viral RNA-dependent RNA polymerase. Transcription requires these proteins and the EBOV VP30 protein (Muhlberger, Lotfering et al., 1998, Muhlberger, Weik et al., 1999).

Filoviral VP30s are zinc- and RNA-binding proteins that are components of the viral nucleocapsid complex (Becker, Rinne et al., 1998, Biedenkopf, Schlereth et al., 2016b, John, Wang et al., 2007, Modrof, Becker et al., 2003, Muhlberger et al., 1999, Nanbo, Watanabe et al., 2013, Schlereth, Grunweller et al., 2016). VP30s are essential for the virus life cycle (Biedenkopf, Lier et al., 2016a, Enterlein, Volchkov et al., 2006, Halfmann, Kim et al., 2008, Muhlberger et al., 1999). A critical role for VP30 is in initiation of viral transcription, a function that is dependent on a stem-loop structure present at the NP gene start site; disruption of the secondary structure leads to VP30-independent transcription (Weik, Modrof et al., 2002). EBOV VP30 also facilitates re-initiation at downstream genes during viral transcription and regulates editing by the viral polymerase during synthesis of nascent mRNAs from the glycoprotein (GP) gene (Martinez, Biedenkopf et al., 2008, Mehedi, Hoenen et al., 2013).

A substantial body of data implicates VP30 phosphorylation as a regulator of its transcriptional function. Dephosphorylation of VP30 N-terminal serine and threonine residues or mutation of these residues to alanine increases pro-transcriptional activity; whereas phosphorylation or mutation to aspartic acid residues inhibits transcription and promotes viral genome replication (Ammosova, Pietzsch et al., 2018, Biedenkopf, Hartlieb et al., 2013, Biedenkopf et al., 2016a, Ilinykh, Tigabu et al., 2014, Kruse, Biedenkopf et al., 2018, Lier, Becker et al., 2017, Martinez, Volchkova et al., 2011, Modrof, Muhlberger et al., 2002, Tigabu, Ramanathan et al., 2018). VP30 interacts with NP, and this influences VP30 phosphorylation levels (Biedenkopf et al., 2013, Kirchdoerfer, Moyer et al., 2016, Kruse et al., 2018, Lier et al., 2017, Xu, Luthra et al., 2017). NP recruits the host PP2A-B56 protein phosphatase through an LxxIxE motif, promoting VP30 dephosphorylation and viral transcription (Kruse et al., 2018). This activity likely explains the importance of VP30-NP interaction for RNA synthesis, as was demonstrated in studies that defined the structure of the VP30-NP interaction interface (Kirchdoerfer et al., 2016, Xu et al., 2017).

Recently, a comprehensive affinity tag-purification mass spectrometry (AP-MS) analysis of the EBOV-host protein-protein interactome identified a number of VP30 interacting host proteins, including retinoblastoma binding protein 6 (RBBP6) (Batra, Hultquist et al., 2018). RBBP6 is a multi-domain protein with E3 ubiquitin ligase activity that has been implicated in cell cycle progression, nucleic acid metabolism, cell proliferation and differentiation (Ntwasa, 2016). RBBP6 possesses a PPxPxY motif that was demonstrated to interact with VP30 at a site that binds a PPxPxY motif on NP. RBBP6 can compete with NP for binding to VP30, thereby inhibiting viral gene expression. In this study, we characterize VP30 interaction with an additional three host proteins that possess PPxPxY motifs. The data demonstrate that the extended sequence PxPPPPxY mediates optimal binding and that proteins sharing this motif compete with NP for binding to a common site on VP30, alter VP30 phosphorylation, modulate viral mRNA transcription and influence viral infectivity. These findings reveal a unique virus-host interaction where multiple host factors target the same viral interface to alter replication efficiency.

## Results

### Multiple host proteins possessing PPxPxY motifs interact with EBOV VP30

A previous study identified RBBP6 as a VP30 interacting protein and demonstrated that a PPxPxY motif mediates the interaction with VP30; this is reminiscent of a PPxPxY motif present in the EBOV NP protein that binds to VP30 (Figure 1A) (Batra et al., 2018). The same study also identified the cellular proteins hnRNP L and hnRNPUL1 in both HEK293T and Huh7 cells and PEG10 in Huh7 cells as VP30 interactors containing the sequence PPxPxY (Figure 1B) (Batra et al., 2018). To further validate these interactions, ectopically expressed FLAG-tagged host proteins were co-expressed with HA-tagged VP30 in HEK293T cells and pull downs were performed using anti-FLAG beads (Figure 1C). All three host proteins interacted robustly with VP30, similar to RBBP6. The interaction between VP30 and endogenous host proteins was further confirmed in HEK293T cells by pull-down of HA-VP30, followed by western blot (Figure 1D). PEG10 is highly expressed in hepatocellular carcinoma cells and was previously identified as an interactor in only Huh7 cells, therefore we confirmed the interaction of VP30 with endogenous PEG10 in these cells (Figure EV1).

**Figure 1.**
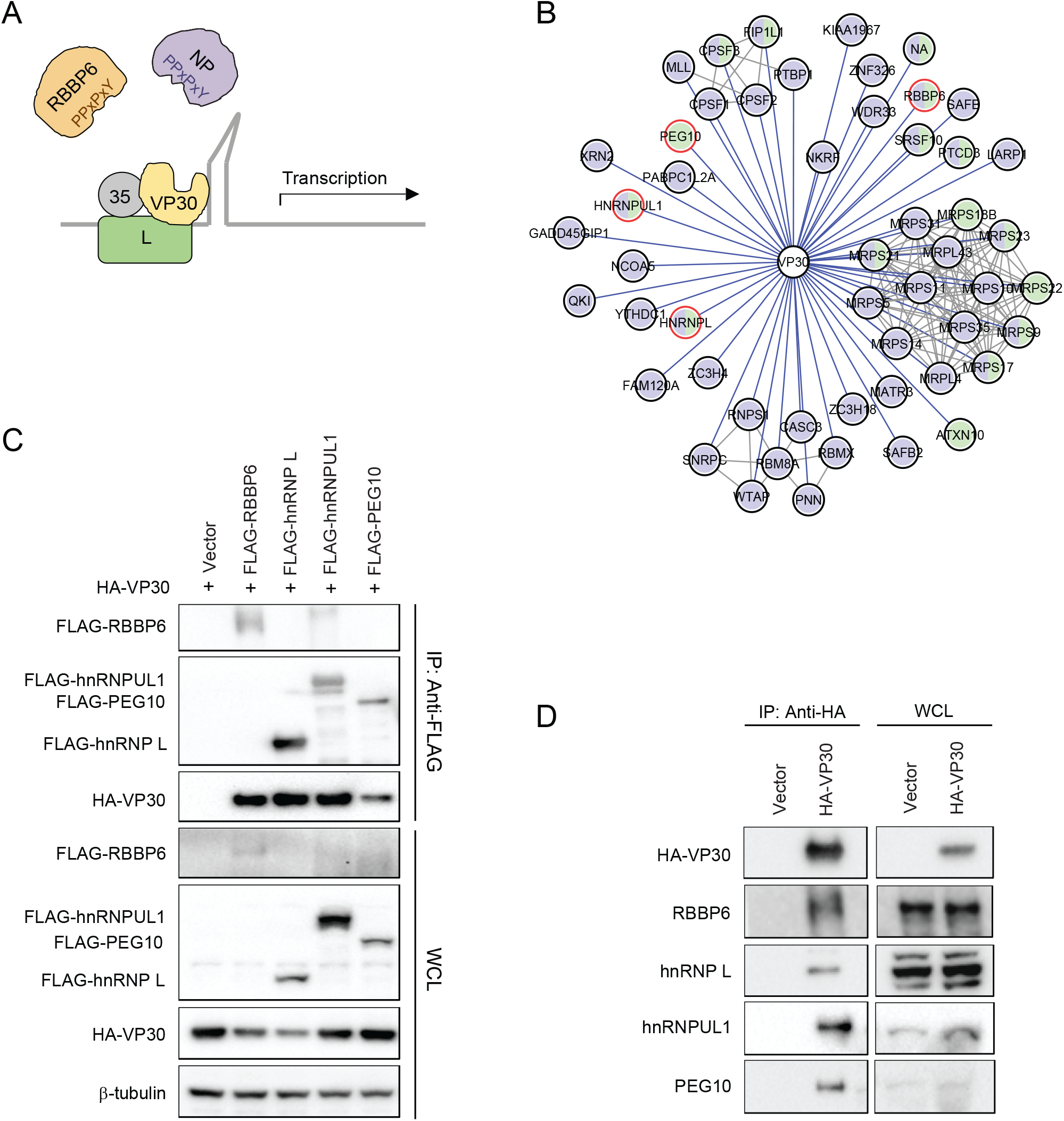
VP30 interacts with multiple host proteins that contain PPxPxY motifs. **(A)** Both EBOV NP and human RBBP6 share a common PPxPxY motif for binding to Ebola virus transcription factor VP30. Interaction between NP and VP30 modulates viral RNA synthesis. RBBP6 outcompetes NP for VP30 binding and inhibits viral infection. **(B)** An EBOV VP30-human protein-protein interaction network. Purple and green prey nodes indicate that the protein was identified as a VP30 interactor in HEK293T and Huh7 cells, respectively; purple-green bifurcation indicates interaction in both cell types. Grey lines correspond to human-human protein-protein interactions curated in the publicly available CORUM database. Host proteins containing PPxPxY motif(s) are highlighted. **(C)** Co-immunoprecipitation (co-IP) between VP30 and the indicated PPxPxY containing host factors. Empty expression plasmid (Vector) or FLAG-tagged host protein expression plasmids were co-transfected with HA-VP30 expression plasmid in HEK293T cells. Anti-FLAG immunoprecipitation (IP: Anti-FLAG) was performed. Western blots of IP and whole cell lysates (WCL) are shown. Anti-β-tubulin was used as a loading control for the WCLs. **(D)** Cells were transfected with empty vector or HA-VP30 expression plasmid. Immunoblotting with anti-HA tag or antibodies to the indicated host proteins was performed after IP with anti-HA antibody (IP: Anti-HA) or on whole cell lysates (WCL).

### Multiple PPxPxY proteins can modulate EBOV RNA synthesis

Based on prior data indicating that RBBP6 inhibits EBOV RNA synthesis, the effect of over-expression of the three host proteins was assessed in an EBOV minigenome (MG) assay. Plasmids expressing hnRNP L, hnRNPUL1, and PEG10 were transfected at two different concentrations along with the plasmids required for the MG assay. Empty vector and RBBP6 were included as controls. MG activity was significantly reduced in hnRNP L- and PEG10-expressing cells (Figure 2A, top). In contrast, hnRNPUL1 increased MG activity. Western blotting of lysates from these experiments demonstrated different degrees of inhibition of viral protein expression at the higher concentration of hnRNP L, hnRNPUL1 and PEG10 plasmids. However, inhibition of viral protein expression was absent at the lower concentrations, where modulation of MG activity was still visible (Figure 2A, bottom). The effect of knocking down endogenous levels of hnRNP L and hnRNPUL1 was also assessed, with RBBP6 siRNA and an irrelevant siRNA serving as controls. 24 h post-siRNA transfection, cells were transfected with the MG assay plasmids. As previously reported, RBBP6 knockdown increased MG assay signal, consistent with RBBP6 acting as an inhibitor of viral RNA synthesis. hnRNP L knockdown also increased the signal (Figure 2B), correlating with it being an inhibitor. hnRNPUL1 knockdown did not significantly affect MG activity (Figure 2B). Because PEG10 protein was difficult to detect in HEK293T cells by western blotting, we carried out a MG assay in Huh7 cells. In these cells, PEG10 knockdown exhibited a modest increase in MG activity (Figure EV2A, B). These data suggest that multiple PPxPxY host proteins can individually exert a negative effect on EBOV RNA synthesis. To address whether these proteins have the capacity to act cooperatively, double knockdown of RBBP6 and hnRNP L was performed in the context of the MG assay. The results demonstrated an additive effect, suggesting that both host proteins can simultaneously modulate viral RNA synthesis (Figure 2C).

**Figure 2.**
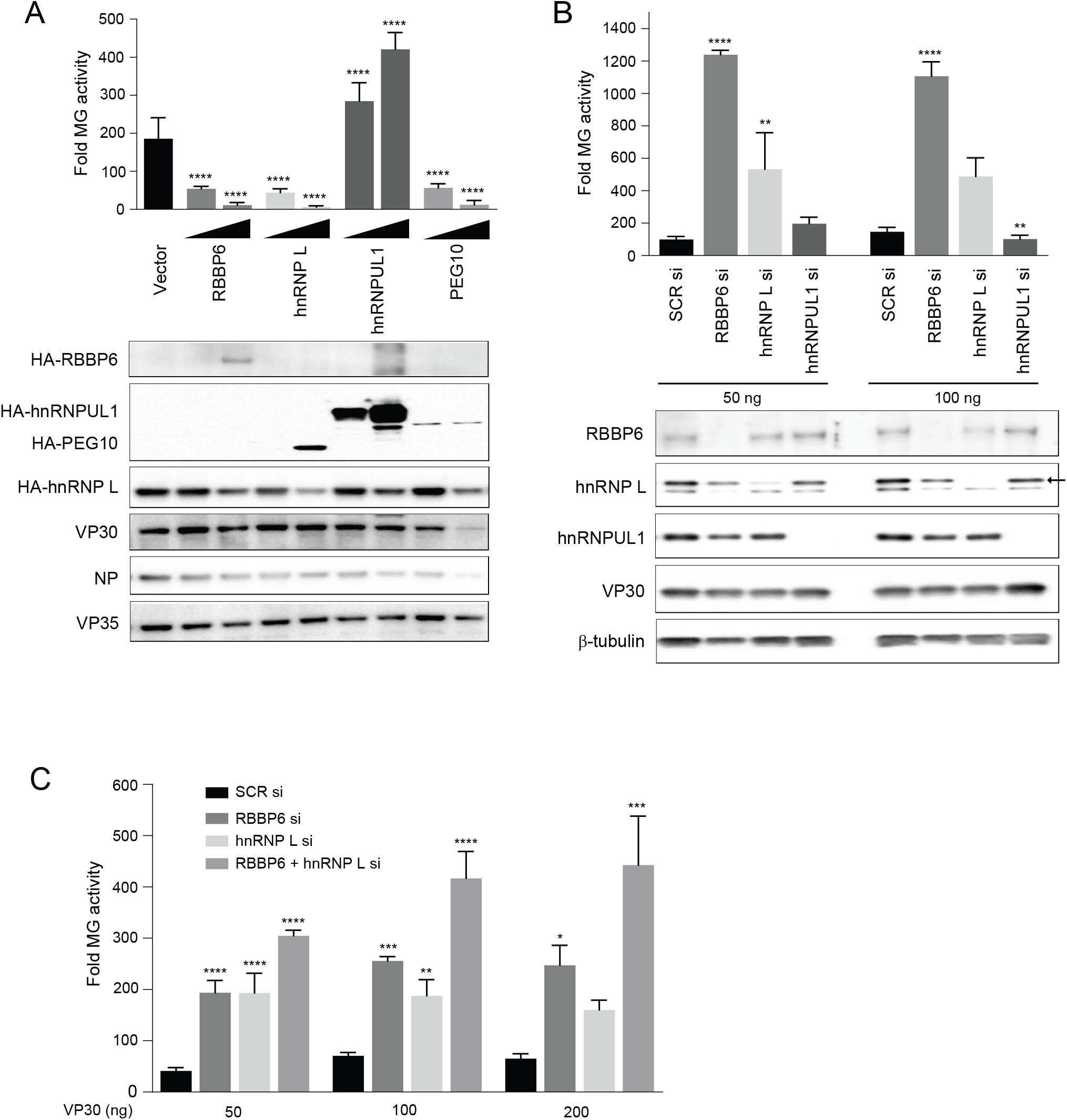
Functional interaction between VP30 and multiple host proteins that contain PPxPxY motifs. **(A)** The effect of over-expression of the indicated host proteins on EBOV RNA synthesis was assessed using an EBOV MG assay. HEK293T cells were transfected with plasmids expressing components of the MG assay and either 50 or 500 ng of expression plasmids for the indicated host factors. Empty expression plasmid (vector) served as a control. Luciferase activity was measured 48h post-transfection and fold MG activity was calculated relative to a no VP30 control. The data represent the mean ± S.D. from one representative experiment (n=3) of at least three independent experiments. Statistical significance was calculated using ANOVA with Tukey’s multiple comparisons test. ****p<0.00005; ***p<0.0005, **p<0.005, *p<0.05. Immunoblots detecting the levels of over-expressed host proteins, VP30 and β-tubulin are shown. **(B)** Minigenome activity upon knockdown of host genes. HEK293T cells were transfected with scrambled siRNA or siRNA targeting RBBP6, hnRNP L or hnRNPUL1. Twenty-four hours post-transfection, cells were transfected with MG assay plasmids. Data represent mean ± S.D. from one representative experiment (n=3) of at least three independent experiments. **(C)** MG activity was assessed as in panel B upon single or double knockdown of RBBP6 and hnRNP L.

### Molecular dissection of PPxPxY-mediated interactions with VP30

There is only one PPxPxY motif within the proline (Pro)-rich region of RBBP6. However, in hnRNP L, PEG10 and hnRNPUL1, two or three copies of this motif are present (Figure 3A, left). The amino acid sequences of peptides derived from host factors, including 10 amino acids flanking each PPxPxY motif, were aligned (Figure 3A, right). Sequences corresponding to each of these extended peptides (labelled as 1-3) were cloned as N-terminal GFP fusions. The fusion proteins were tested in co-immunoprecipitation assays for their ability to bind VP30. In each case, only one of the tested peptides from hnRNP L (hnRNP L_2), hnRNPUL1 (hnRNPUL1_3) and PEG10 (PEG10_2) strongly interacted with VP30, suggesting that the sequence context of the motif plays a significant role in binding (Figure 3B). Also observed was a weak interaction between hnRNPUL1 peptide 1 and VP30.

**Figure 3.**
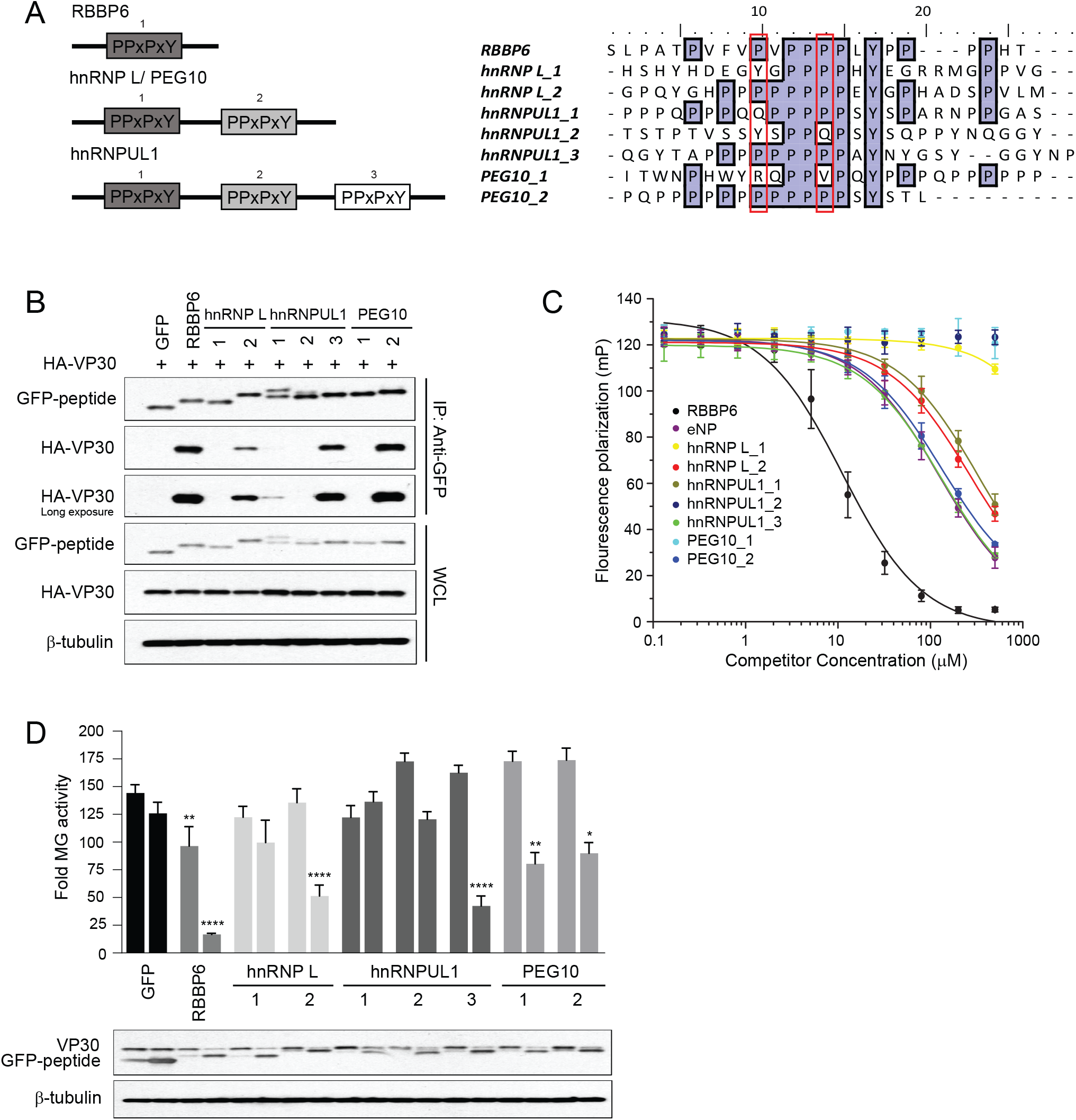
PPxPxY containing peptides derived from host proteins interact with VP30 and modulate EBOV RNA synthesis. **(A)** Schematic representation of the arrangement and number of PPxPxY motifs present in the indicated host proteins (left panel). Multiple sequences aligned using ClustalW, of peptides containing PPxPxY motif derived from VP30 interacting host proteins (right panel). **(B)** Co-IP between HA-VP30 and either GFP or GFP-fused to peptides derived from the host proteins was performed. Western blots of the co-IP and whole-cell lysate (WCL) are shown. **(C)** Equilibrium dissociation curves of FITC-RBBP6 from eVP30 (130-272) as it is outcompeted by increasing concentrations (0.13-500 μM) of PPxPxY containing peptides. Fluorescence polarization was determined with constant concentrations of FITC-RBBP6 and eVP30 (130-272), at 0.50 μM and 3.8 μM respectively. Experiments were performed in two independent duplicates. Error bars represent standard deviation. **(D)** Minigenome activity upon transient expression of GFP or GFP-fused PPxPxY-containing peptides. Plasmids expressing GFP fused peptides were transfected at 50 and 500 ng amounts Representative immunoblots of the lysates detecting expression of the GFP-peptide fusions, VP30 and β-tubulin are shown. Data represent the mean ± S.D. from one representative experiment (n=3) of at least three independent experiments. Statistical significance was calculated relative to the GFP control for each concentration.

To quantify the strength of individual peptide binding, and to determine if the peptides can target the VP30 interface bound by RBBP6 and NP, a fluorescence polarization (FP) assay was employed in which a purified C-terminal domain of EBOV VP30 (eVP30_130-272_) and a PPxPxY containing peptide derived from EBOV NP were used (Figure 3C). The RBBP6 peptide displayed the highest binding affinity. hnRNPUL1_3 and PEG10_2 demonstrated affinities similar to the NP peptide, whereas hnRNP L_2 and hnRNPUL1_1 exhibited slightly lower affinities. hnRNP L_1, hnRNP UL1_2 and PEG10_1 exhibited minimal to no binding in either the FP or pulldown assays. Overall, these data support the extended sequence PxPPPPxY as contributing to optimal binding; however, the sequence composition surrounding this motif contributes to the strength of interaction.

We compared the activity of the single PPxPxY-motif peptide fusions from hnRNP L, hnRNPUL1, and PEG10 to that of the GFP-RBBP6 peptide in the MG assay (Figure 3D). RBBP6 peptide was inhibitory, consistent with prior studies (Batra et al., 2018). Each of the remaining peptides that interacted in the co-precipitation and FP assays inhibited MG activity, including hnRNPUL1 peptide 3, despite full-length hnRNPUL1 acting as a MG enhancer (Figure 3D, EV3A).

The EBOV protein VP40 possesses overlapping PTAP and PPEY motifs (PTAPPEY) that function as late domains in viral budding (Licata, Simpson-Holley et al., 2003). To determine whether this sequence might mediate an interaction with VP30 and to further address the specificity of Pro-rich motifs to interact with VP30, EBOV VP40/VP30 co-IPs were performed. VP40 did not co-precipitate with VP30 (Figure EV3B). Furthermore, fusion of the N-terminal VP40 PTAPPEY sequence to GFP also did not facilitate co-precipitation of VP30. Mutation of the late domain motif to contain five proline residues (PPPPPEY) failed to confer interaction. However, mutation of the VP40 peptide to include an additional sixth proline (PPPPPPEY) conferred interaction (Figure EV3C). These data further support the extended PxPPPPxY motif as conferring efficient binding to VP30.

We also tested an RBBP6 peptide (bRBBP6 peptide) derived from a fruit bat (*Rousettus aegyptiacus*) that serves as MARV reservoir hosts in co-precipitation assays with VP30. bRBBP6 peptide differs at only two positions from the human RBBP6 peptide (hRBBP6 peptide) and binds robustly to VP30 with a near identical affinity, which further substantiates the requirement of the PxPPPPxY motif for optimal binding to VP30 (Figure EV3D, E). When titrated in the MG assay, bRBBP6 was modestly less inhibitory to MG activity, as compared to the hRBBP6 peptide (Figure EV3F).

The binding in solution between VP30 and different host peptides, including the interacting hRBBP6, hnRNPUL1_1, hnRNPUL1_3, hnRNPL_2, and PEG10_2, and the non-interacting hnRNPUL1_2, was further characterized with hydrogen deuterium exchange (HDX) mass spectrometry. VP30 binds nearly identically to RBBP6, hnRNPUL1_1, hnRNPUL1_3, hnRNP L_2 and PEG10_2 peptides as seen in the HDX kinetic curves, indicating that each of the peptides binds to a similar region on VP30 (Figure 4A and Figure EV4A). Statistical analysis of deuterium uptake differences identified two strong peptide binding regions on VP30 from residue L189 to V210 and from residue A224 to D231. The two regions contain seven and four binding residues, respectively, that were identified by X-ray crystallography (Batra et al., 2018), demonstrating good consistency of the two approaches and the ability of HDX to give an accurate picture of the binding. The middle region from residue Y211 to E223 presents as a weaker binding region where the differences in HDX are much smaller between bound and unbound, also consistent with the X-ray structure that show binding via only one residue, R213. Although there is clear binding from residues 224-231, the region 230-237 shows different character in the kinetics of HDX. Its extent for the bound state converges at long times with that of the unbound, indicating a larger off rate and weaker binding at the end of the peptide ligand. We further mapped the HDX differences at 4 h onto the VP30 crystal structure (PDB:6E5X), which illustrates common binding regions among different host factor peptides (Figure 4B).

**Figure 4.**
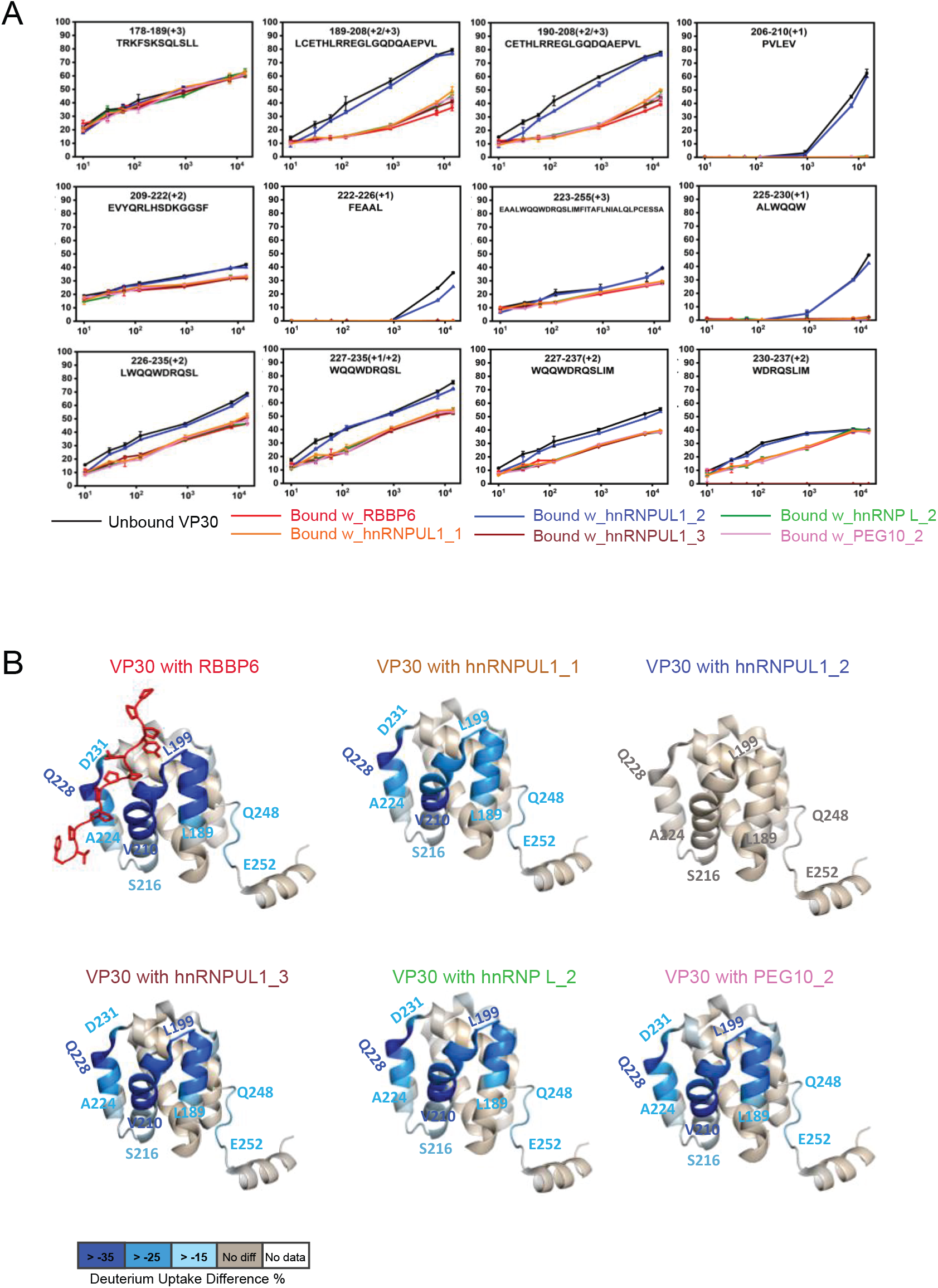
HDX-MS analysis of VP30 and host factor peptide interactions. **(A)** HDX kinetics data for indicated host peptides incubated with VP30 or VP30 alone are shown. VP30 peptide amino acid residue positions and sequences are indicated at the top of each curve. **(B)** Deuterium uptake differences were mapped onto the VP30 structure (PDB: 6E5X), revealing similar binding regions for the peptides. Residue-level uptake difference was achieved by overlapping peptides.

In addition, the HDX studies revealed another region, residues Q248 to E252, that either describes tightening dynamics or undergoes remote conformational changes upon peptide binding (Figure 4B, EV4B). For hnRNPUL_2, differential HDX kinetic analysis (in absence and presence of the ligand) yielded similar HDX, consistent with the lack of its binding in the pull-down and fluorescence polarization assays.

To determine whether each of the full-length host proteins target the NP/RBBP6 binding site on VP30, VP30-host protein interaction was assessed in the presence of increasing amounts of NP (Figure 5A). As NP levels increased, VP30 interaction with hnRNP L, hnRNPUL1 and PEG10 was disrupted. As previously reported, increasing amounts of NP could also disrupt the VP30-RBBP6 interaction (Batra et al., 2018). Select point mutations within the NP binding cleft on VP30, including E197A, W230A, or combinations of mutations that included these residues, were previously demonstrated to disrupt binding to both NP and RBBP6 (Batra et al., 2018; Xu et al. 2017; Kirchdoerfer et al., 2016). Similarly, E197A, W230A, or combinations of mutations that included these residues exhibited loss of binding to hnRNP L, hnRNPUL1, and PEG10 (Figure 5B, EV5A,B). These data further support an interaction at the same site on VP30, as well as similar modes of binding.

**Figure 5.**
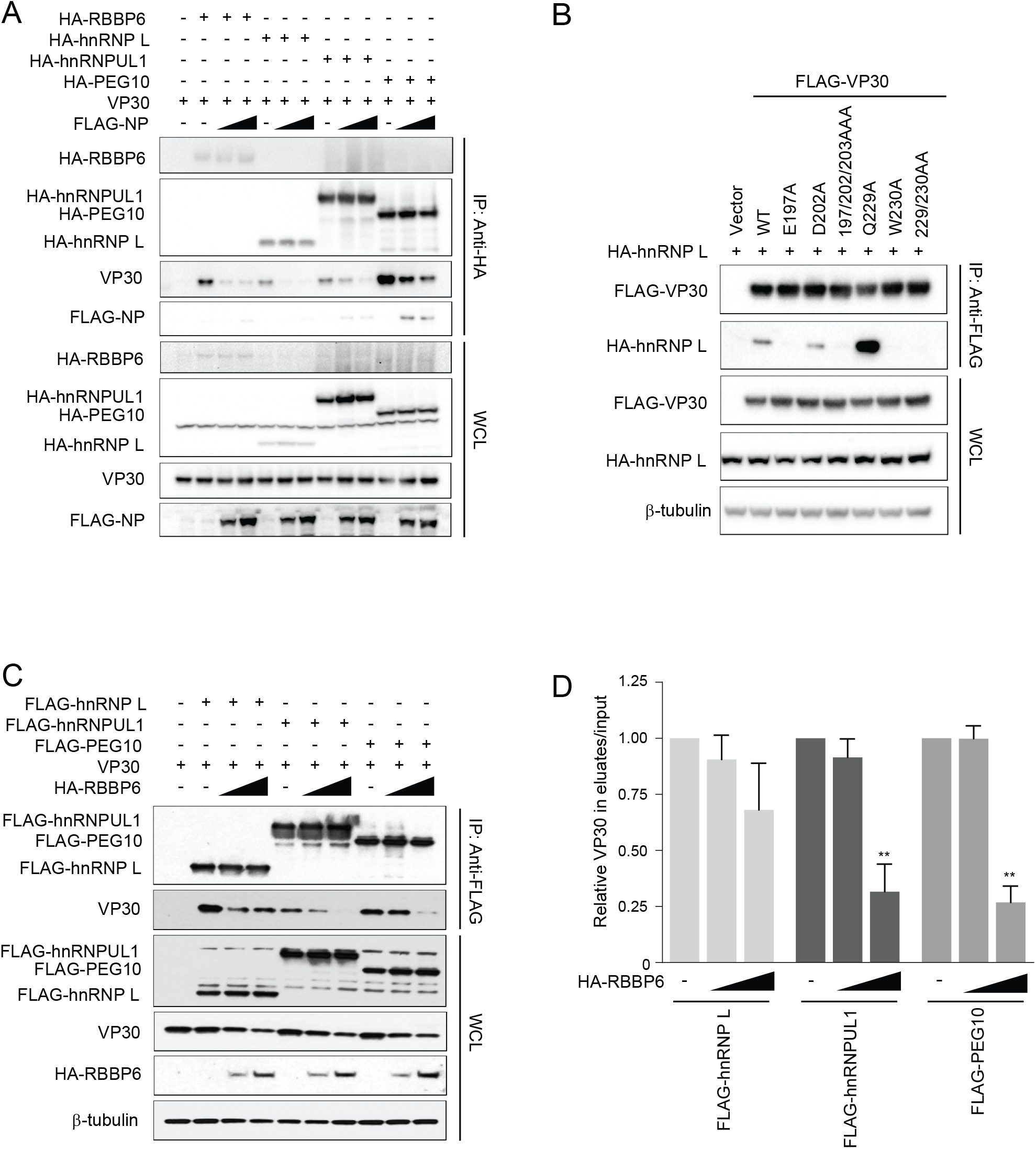
PPxPxY containing host proteins target the NP binding cleft on VP30. **(A)** Co-IP experiment demonstrating that NP interferes with VP30-PPxPxY binding. VP30 was co-expressed with the indicated HA-tagged PPxPxY proteins and either empty vector (-) or increasing amounts of FLAG-tagged NP expression plasmid. IPs were performed with anti-HA antibodies followed by western blots of IPs or WCL by using anti-HA, anti-FLAG and anti-VP30 antibodies. β-tubulin served as a loading control. **(B)** Representative immunoblots of co-IPs between hnRNP L and the indicated VP30 mutants previously described to exhibit varying degrees of NP binding activity. **(C)** Co-IP between VP30 and indicated host proteins upon titration of RBBP6. VP30 was co-expressed with FLAG-tagged PPxPxY containing host proteins and either a 5-fold or 10-fold excess of RBBP6 plasmid. IP was performed using anti-FLAG beads followed by immunoblotting using the indicated antibodies. **(D)** Densitometric analysis of the western blots shown in panel C. The graph represents relative amount of VP30 present in the eluates compared to the input.

If there is a shared binding site on VP30, it would be expected that different PxPPPPxY motif containing proteins could compete with each other for binding with VP30. To test this possibility, we performed competition co-IPs between VP30 and FLAG-tagged hnRNP L, hnRNPUL1 or PEG10 in the presence of HA-tagged RBBP6. Western blotting and densitometric analysis of VP30 levels in eluates versus input suggested that increasing amounts of RBBP6 significantly disrupts the interaction between VP30 and hnRNPUL1 or PEG10 (Figure 5C, D). Competition with hnRNP L was less effective, but as RBBP6 levels increased, hnRNP L binding was modestly reduced. These data further suggest that multiple host proteins may compete with each other in infected cells.

### Residues critical for VP30 interaction and MG assay inhibition

To further define the critical residues within the extended PxPPPPxY motif required for binding to VP30, we performed alanine scanning mutagenesis across the proline and tyrosine residues in the RBBP6 peptide, testing the VP30 binding activity of these mutants in a co-IP assay. Mutation of the motif (P_1_xP_2_P_3_P_4_P_5_xY) at prolines 2 and 5, and the tyrosine greatly impaired binding (Figure 6A). The critical role of these residues was further supported by FP analysis with wild-type and mutated RBBP6-derived peptides (Figure 6B). GFP fusion peptides with alanine substitution mutations at multiple positions including prolines 1, 3 and 4 resulted in a weaker interaction with VP30 compared to the wildtype (WT) peptide (Figure 6C). When the mutant peptides were analyzed for their ability to inhibit the MG assay as GFP fusions, mutation of the proline residue at position 5 or the tyrosine lost inhibitory activity as compared to the WT RBBP6 peptide. Surprisingly, the P2A mutant retained inhibitory activity despite absence of detectable interaction (Figure 6D), suggesting a potential off-target effect.

**Figure 6.**
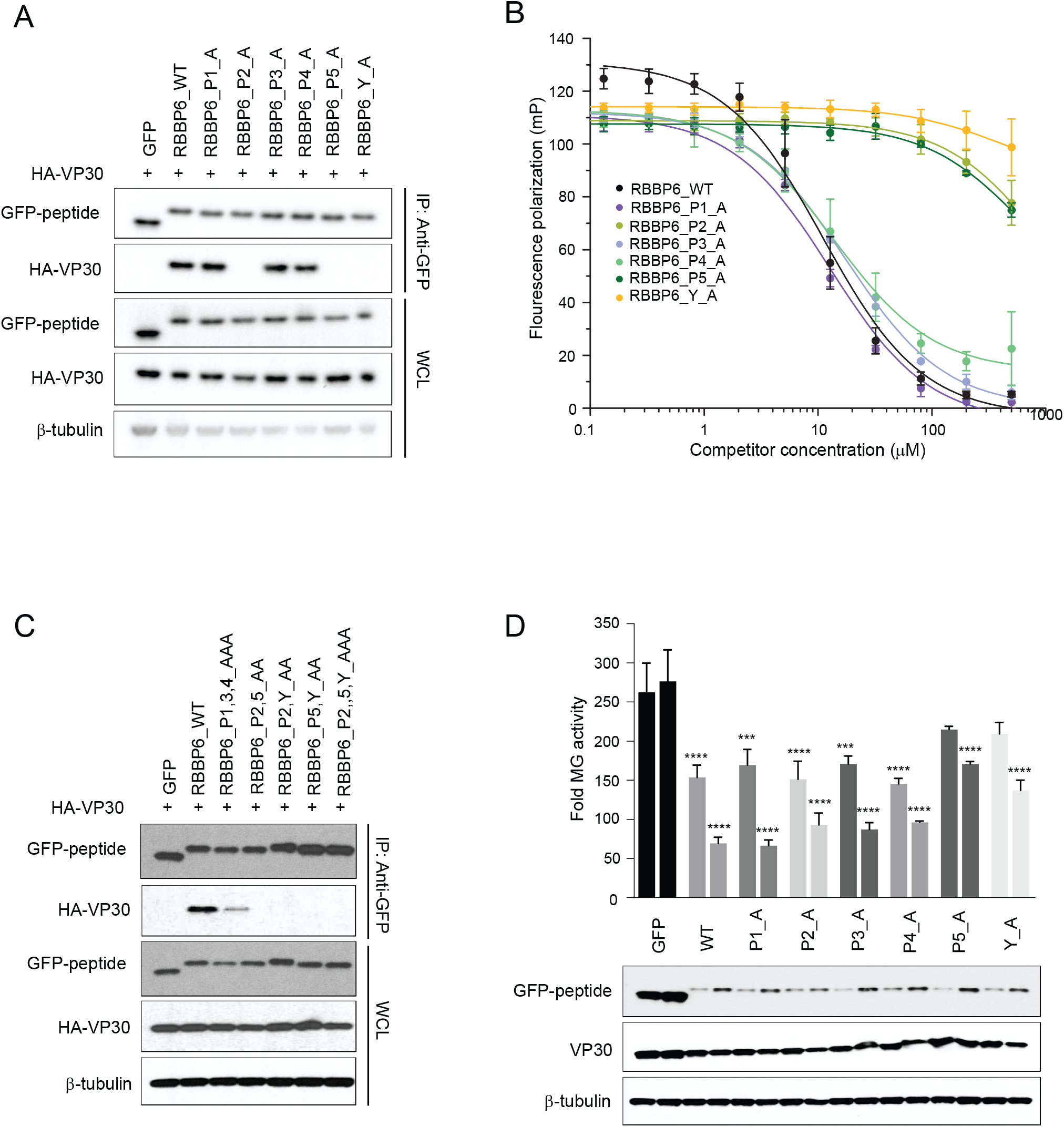
Defining the contribution of PPxPxY residues to VP30 interaction. **(A)** Co-IP between HA-VP30 and either GFP, or GFP-fused to WT or mutant RBBP6 peptides. Each proline or tyrosine residue in the P_1_xP_2_P_3_P_4_P_5_xY motif was mutated to alanine and the indicated mutant peptides were expressed as GFP fusions. Co-IPs of HA-VP30 with GFP or GFP-peptide fusions were performed using anti-GFP magnetic beads (IP:Anti-GFP). Whole cell lysates (WCL) were blotted with anti-GFP, anti-HA and anti-β-tubulin antibodies. **(B)** Equilibrium dissociation curves of FITC-RBBP6 from eVP30 (130-272) as it is outcompeted by increasing concentrations (0.13-500 μM) of RBBP6 alanine mutants. Fluorescence polarization was determined with constant concentrations of FITC-RBBP6 and eVP30 (130-272) at 0.50 μM and 3.8 μM respectively. Experiments were performed in two independent duplicates. Error bars represent standard deviation. **(C)** Co-IP between HA-VP30 and RBBP6 peptide mutants. **(D)** Minigenome activity upon titration of plasmids encoding GFP-RBBP6 peptides with wild-type or single point mutant peptides. GFP alone or GFP-RBBP6 peptide plasmids were transfected at 50 and 100 ng amounts along with the plasmids of the minigenome system. ****p<0.00005; ***p<0.0005, **p<0.005, *p<0.05

### PxPPPPxY motif containing host proteins modulate virus infection

To validate the impact of the host proteins on EBOV replication, virus infections were performed in the context of siRNA knockdown. HeLa cells were transfected with siRNAs against human hnRNP L and hnRNPUL1, alongside control scrambled siRNA or siRNA targeting critical entry factor NPC1, as a positive control, followed by infection with EBOV-EGFP. Twenty-four hours post-infection, cells were fixed and quantified for infection by microscopy. Knockdown of hnRNP L strongly enhanced virus infection, whereas hnRNPUL1 siRNA-treatment was inhibitory, consistent with MG data (Figure 7A,B). Knockdown of NPC1, a host protein essential for virus entry, strongly reduced virus replication, as expected (Carette, Raaben et al., 2011). The efficacy of siRNA-mediated knockdown was confirmed by western blotting using specific antibodies (Figure S1A,B).

**Figure 7.**
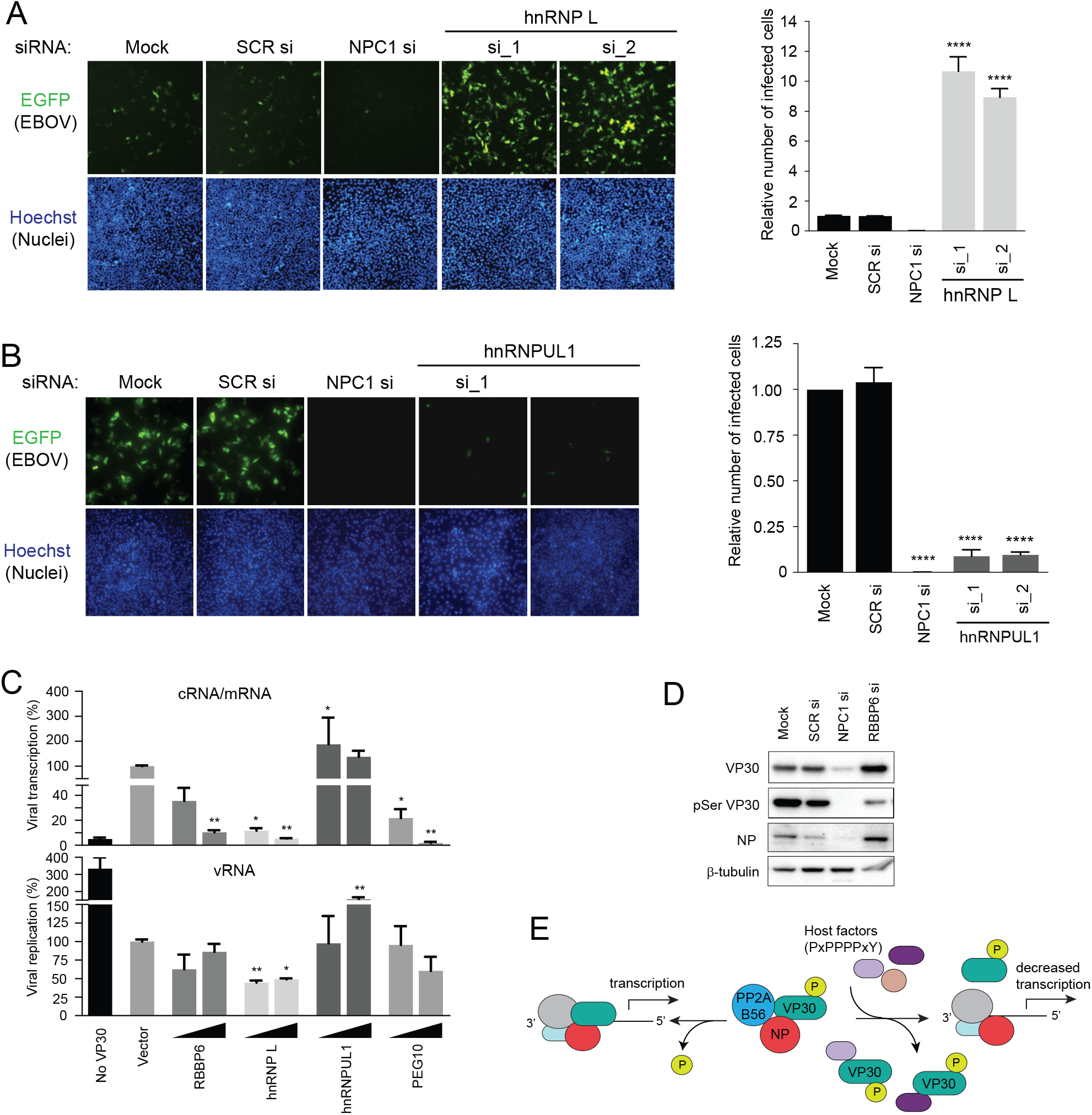
PPxPxY containing host proteins modulate EBOV infectivity, viral transcription and VP30 phosphorylation. **(A)** HeLa cells were mock transfected or transfected with scrambled siRNA (SCR si) or siRNAs targeting NPC1 or hnRNP L (hnRNP L si_1 and si_2), 48 h post-transfection, cells were challenged with EBOV-EGFP at an MOI of 0.05. Twenty-four hours post-infection, cells were fixed, stained for nuclei with Hoechst dye and imaged. The graph shown (right) depicts percent infection as determined by calculating the ratio of number of infected cells to nuclei and shown relative to mock infected cells. Statistical significance was calculated relative to the values obtained in SCR siRNA-treated cells using ANOVA with Tukey’s multiple comparisons test. **(B)** HeLa cells were mock transfected or transfected with scrambled siRNA (SCR si) or siRNAs targeting NPC1 or hnRNPUL1 (hnRNPUL1 si_1 and si_2). After 48 h, cells were infected with EBOV-EGFP at an MOI of 0.5 for 24 h. Hoechst-stained images (left) and representative graphs indicating percent infection (right) are shown. **(C)** Levels of transcription and replication in the MG assay upon over-expression of the indicated host factors. MG assays were performed with 50 and 500 ng of host protein expression plasmid. RNA was isolated and analyzed by qRT-PCR to measure the levels of viral mRNA/cRNA and vRNA. The vector control sample was set to 100% and mean ± SD values from two independent experiments are shown. ***p<0.00005; ***p<0.0005, **p<0.005, *p<0.05. **(D)** Immunoblots to assess the levels of VP30, pSer29 VP30 and NP in EBOV infected lysates upon knockdown of NPC1 or RBBP6. HeLa cells were mock transfected or transfected with siRNAs targeting NPC1 or RBBP6. 48 h post-transfection cells were infected with EBOV at MOI=0.1. At 24 h post-infection, cells were lysed in TRIzol. TRIzol extracted protein fractions were analyzed by western blotting to determine levels of VP30, pSer29 VP30 and NP. **(E)** Proposed model for inhibition of EBOV replication by host factors containing PxPPPPxY motifs. Host proteins present at varying endogenous levels disrupt VP30-NP interaction thereby preventing VP30 dephosphorylation by NP-recruited host phosphatase PP2A-B56, and therefore inhibit virus transcription.

### PxPPPPxY motif containing proteins influence phosphorylation of VP30

VP30 undergoes dynamic phosphorylation by serine-arginine protein kinases (SRPK1 and SRPK2) that regulate its transcription-promoting activity (Biedenkopf et al., 2016a, Lier et al., 2017, Takamatsu, Krahling et al., 2020). NP then recruits a host protein phosphatase, protein phosphatase 2A B56 (PP2A B56), resulting in dephosphorylation of NP-bound VP30. The resulting dephoshorylated VP30 promotes viral transcription (Kruse et al, 2018). VP30-interacting host proteins may prevent the dephosphorylation of VP30 by competing with NP and inhibiting transcription. In contrast, elimination of competing host factors might increase VP30 dephosphorylation and enhance viral transcription. To test this, we measured the amount of viral mRNA/antigenomic RNA (cRNA) and viral genomic RNA (vRNA) in the context of the MG following expression of each host protein. Viral transcription was significantly reduced when RBBP6, hnRNP L or PEG10 were expressed, whereas the presence of hnRNPUL1 resulted in increased viral transcription (Figure 7C). These findings parallel the effect of these proteins on the MG assay (Figure 2A). To further substantiate these findings, we utilized an antibody that specifically recognizes VP30 phosphorylated at Ser29 (Lier et al., 2017; Kruse et al., 2018). siRNA-mediated knockdown of RBBP6 resulted in strongly reduced levels of VP30 phosphorylation in infected cells (Figure 7D). These data suggest a model whereby RBBP6 and other proteins containing PxPPPPxY motifs can disrupt VP30-NP interaction, preventing VP30 dephosphorylation by PP2A B56 and modulating viral transcription (Figure 7E).

## Discussion

This study characterized three novel EBOV VP30-host protein-protein interactions (PPIs) and demonstrated their functional consequences. Each of the interactions involved a PPxPxY motif present in the host protein. hnRNP L, hnRNPUL1, and PEG10 were identified in a recent AP-MS study that defined an EBOV-host PPI network for six EBOV proteins. The network included 194 high-confidence EBOV-human PPIs, among which was the previously characterized VP30 interacting PPxPxY-containing protein RBBP6 (Batra et al., 2018). Sequence analysis of the VP30 binding peptides derived from RBBP6 and NP identified a common PPxPxY motif. Notably, this motif is conserved among NPs of the *Ebolavirus*, *Marburgvirus*, C*uevavirus* and *Dianlovirus* genera within the filovirus family. Based on the RBBP6 data, the VP30 interaction network was inspected for additional proteins possessing PPxPxY motif(s), identifying hnRNP L and PEG10 as having two such motifs and hnRNPUL1 as having three. Co-immunoprecipitation studies confirmed the interaction between VP30 and RBBP6, hnRNP L, hnRNPUL1, and PEG10. Several lines of evidence suggest that these proteins target the same site on VP30. Competition between the PPxPxY proteins and NP for VP30 binding was demonstrated in co-IP studies. Mutations in VP30 that alter NP binding similarly affect binding to each of the host factors; and competition between the host factors for binding to VP30 could also be demonstrated. Further, an FP assay that measured the capacity of host protein-derived peptides to disrupt interaction between the C-terminal domain of VP30 and NP demonstrated that the four host factors target the NP binding site on VP30. We also mapped the effect of peptide binding on the conformational dynamics of VP30 protein by employing HDX-MS. The distinct patterns of HDX suggested that binding of peptides containing an extended PxPPPPxY motif induce significant changes even at the longer time scales, supporting a conformational change in a nearby region (Q248 to E252). All these findings point to a remarkable circumstance where multiple host proteins target a critical virus-virus PPI leading to viral RNA synthesis modulation.

A PPxPxY motif characterizes each of the VP30 interacting proteins. Proline-rich motifs are frequently involved in protein-protein interactions. Domains that bind proline-rich motifs include Src-homology 3 (SH3), WW, EVH1, GYF (also known as CD2-binding domains), UEV domain, and single-domain profilin proteins (Reviewed in (Ball, Kuhne et al., 2005)). Structurally, the EBOV VP30 C-terminal domain is distinct from these previously described proline-rich motif binding domains in that it is largely α-helical in nature (Kirchdoerfer et al., 2016, Xu et al., 2017). Further, co-crystals of the VP30 C-terminal domain and the NP-derived PPxPxY peptide or the RBBP6 peptide demonstrated that the PPxPxY motif does not form the type II helix typical of PPxP motifs (Batra et al., 2018, Kirchdoerfer et al., 2016, Xu et al., 2017). Rather, in the VP30-NP peptide structures the NP residues are in an extended conformation, making multiple contacts with VP30 residues (Xu et al., 2017). These unique features make understanding the VP30-PPxPxY interaction of particular interest.

The identification of several host proteins that interact with VP30 via PPxPxY motifs allowed us to characterize the basis of this unique Pro-rich motif interaction. To better understand the specificity of different PPxPxY motifs for VP30, pulldowns were performed using peptide fusions to GFP in which each of the individual motifs was flanked on each side by ten amino acid residues. These experiments demonstrated differential binding activity. For those proteins with two or three PPxPxY motifs, in each case only a single motif robustly interacted with VP30 (motif 2 for hnRNP L, motif 3 for hnRNPUL1 and motif 2 for PEG10). FP assay-based measurements of the equivalent peptides yielded a comparable pattern of interactions and demonstrated a range of binding affinities. The RBBP6 peptide was the tightest binder. hnRNPUL1 motif 3 and PEG10 motif 2 behaved similarly to each other and to the PPxPxY peptide derived from eNP. The hnRNP L motif 2 and hnRNPUL1 motif 1 exhibited comparable binding affinities to each other, consistent with the results of the co-IP assays, in which hnRNP L motif 2 bound weakly and hnRNPUL1 motif 1 yielded a barely detectable interaction. By aligning the various motifs and ranking them based on affinities, PxPPPPxY correlated with a stronger interaction. Even among those peptides containing this motif variations were seen, suggesting that diversity in the “x” positions or flanking sequences may also influence binding. Contributions of nearby residues to Pro-rich motif binding has been demonstrated in other contexts as well (Ball et al., 2005).

Because the RBBP6 peptide consistently showed robust binding to VP30 and the highest inhibitory activity in MG assays, the contribution of individual residues to binding was assessed by mutagenesis. The data support critical roles for the second and fifth proline residues and the tyrosine residue, with each individual mutation abrogating the interaction when assessed either by co-IP or FP assay. Interestingly, the P2A mutant retained the capacity to inhibit viral RNA synthesis in the minigenome assay, despite its reduced binding in co-IP assays. This latter finding could reflect a weaker interaction with VP30 that is nonetheless sufficient to inhibit MG activity. Alternatively, there may be off-target effects of expressing the peptide fused with GFP that impact viral RNA synthesis. To address specificity of proline-tyrosine motifs to interact with VP30, a set of experiments were performed with the EBOV matrix protein VP40. EBOV VP40 possess two overlapping late domain motifs _7_PTAPPEY_13_ resembling the PPxPxY motif. The PPxY late domain motif mediates interactions with WW domain host proteins that facilitate VP40 budding and release from cells (Han, Sagum et al., 2016, Han, Sagum et al., 2017, Harty, Brown et al., 2000, Liang, Sagum et al., 2017, Martin-Serrano, Eastman et al., 2005, Timmins, Schoehn et al., 2003, Yasuda, Nakao et al., 2003). Despite the similarity of the VP40 proline-tyrosine motif to PPxPxY, an interaction between VP30 and VP40 was not detected, either with full-length VP40 or when a peptide encompassing the late domain motif of VP40 was fused to GFP. This indicates that VP30 does not bind typical WW domain ligands. Mutation of the VP40 peptide to match the PxPPPPxY motif did confer weak binding, further supporting the conclusion that this motif confers binding. However, the relatively weak interaction with this mutated VP40 peptide provides further evidence that sequences in the vicinity of the motif influence binding. Cumulatively, these studies further define the VP30 interface as a novel proline-tyrosine interacting motif and provide a foundation for further elucidation of its structural basis and functional significance.

That the interactions of VP30 with the endogenous host proteins are functionally significant is supported by the demonstrations that modulation of RBBP6, hnRNP L, PEG10, and hnRNPUL1 levels regulate EBOV RNA synthesis, as measured by minigenome and infection assays. In support of their role as cellular modulators of EBOV replication, over-expression of full-length RBBP6, hnRNP L and PEG10 inhibited EBOV RNA synthesis, whereas transfection of full-length hnRNPUL1 enhanced MG activity. Similarly, expression of interacting PPxPxY-containing peptides also inhibited RNA synthesis. That hnRNPUL1 enhances activity whereas hnRNP L, PEG10 and the GFP-peptide fusions, including GFP-hnRNPUL1 peptide 3 inhibit MG activity, is notable. Interestingly, a mutant of VP30 within the VP30-NP/RBBP6 binding interface, VP30_E197A/D202A/Q203A_, confers enhanced signal in the MG assay (Xu et al., 2017). A possible explanation for this phenomenon would be that binding affinity between VP30 and NP has evolved to confer less than maximal transcription and, when the VP30/NP affinity is reduced to a particular degree, as in the case of VP30_E197A/D202A/Q203A_ or when a competitor with affinity for VP30 is present at the appropriate concentration, activity is enhanced. According to this model, the discrepancy in functional outcomes between expression of hnRNPUL1 versus GFP-hnRNPUL1 peptide 3 would be explained if the GFP-peptide fusion has higher affinity for VP30 or, if the GFP-peptide fusion expresses to higher levels.

Given that multiple host proteins can target the NP binding site on VP30, it was of interest to determine whether, in cells, more than one host protein can simultaneously impact VP30 function. We therefore asked whether knockdown of the two host factors with the largest individual effect on EBOV RNA synthesis would result in additive effects. The minigenome assays performed suggested that this was the case. How several different host factors together modulate EBOV gene expression will require additional study. However, relative expression levels and localizations in the cell will be expected to influence the outcome.

A substantial body of literature implicates VP30 phosphorylation as a regulator of its transcriptional functions (Ammosova et al., 2018, Biedenkopf et al., 2013, Biedenkopf et al., 2016a, Ilinykh et al., 2014, Kruse et al., 2018, Lier et al., 2017, Martinez et al., 2011, Modrof et al., 2002, Tigabu et al., 2018). Further, NP recruits a cellular protein phosphatase to dephosphorylate NP-bound VP30 (Kruse et al., 2018). Previous data demonstrated the capacity of RBBP6 to target the NP binding site on VP30 and to impair viral RNA synthesis and infectivity (Batra et al., 2018). Data in this study demonstrating modulation of viral RNA synthesis by hnRNP L, hnRNPUL1 and PEG10, suggested a model whereby disruption of the NP-VP30 interaction by a host factor would alter NP-mediated VP30 dephosphorylation. If levels of phospho-VP30 rise as a consequence, this would be expected to inhibit virus transcription and gene expression. Fitting this model, expression of RBBP6, hnRNP L, or PEG10 inhibited transcription and hnRNPUL1 enhanced transcription in the MG assay. In each case, effects on mRNA synthesis were more dramatic than effects on products of replication. Further, knockdown of endogenous RBBP6 decreased VP30 phosphorylation levels and increased viral gene expression. Therefore, competition for VP30-NP binding appears to modulate viral transcription by modulating VP30 phosphorylation levels.

It is striking that EBOV maintains a VP30-NP interface critical for replication that can be directly targeted by multiple host factors. While RBBP6, hnRNP L and PEG10 appear to act as restriction factors, in that they prevent VP30 dephosphorylation, inhibiting viral gene expression, it is also possible that EBOV takes advantage of these interactions as a regulatory mechanism. For example, early in infection, when NP levels are relatively low, the presence of competitors for binding to VP30 could reduce dephosphorylation to dampen viral transcription, perhaps as a means to delay recognition by the immune system. Given that our data suggests cooperation between multiple host factors that inhibit viral transcription, it will be of interest to identify cell types with different levels of the host PPxPxY proteins to determine how this shapes the replication cycle.

In addition, these data do not preclude effects of VP30 on host protein function. RBBP6, hnRNP L and hnRNPUL1 are each RNA binding proteins with a number of biological functions, including the regulation of various RNA processing activities, RNA degradation, pre-mRNA splicing and mRNA translation. Beyond this are effects on cellular proliferation, dsDNA break repair and other processes, such as, for hnRNP L, on interferon-β gene expression (Cao, Liu et al., 2019, Gurunathan, Yu et al., 2015, Hong, Jiang et al., 2013, Peng, Zhu et al., 2017, Seo, Kim et al., 2017, Zhou, Li et al., 2017). PEG10 is derived from a retrotransposon gag-pol gene (Xie, Pan et al., 2018). Its mRNA contains two open reading frames (ORFs) with the first ORF encoding a gag-like protein and the second encoding a pol-like protein. A -1 frameshift occurs during translation leading to production of a gag-pol fusion protein. PEG10 is highly expressed in the placenta, adrenal gland, ovary, testis and brain. It plays roles in placenta formation and adipocyte differentiation and its deletion in mice is embryonic lethal (Hishida, Naito et al., 2007, Xie et al., 2018). PEG10 is also upregulated in cancer where it plays a role in promoting cellular proliferation. It remains to be determined whether VP30 interaction modulates the functions of these host proteins and what impact this might have on virus infection.

## MATERIALS AND METHODS

### Cell lines

HEK293T (human embryonic kidney SV40 Tag transformed; ATCC, CRL-3216), Huh7 (generous gift from the Gordan lab at UCSF) and HeLa cells (ATCC, CCL-2) were maintained in Dulbecco’s modified Eagle’s medium (Corning Cellgro or ThermoFisher scientific) with 10% fetal calf serum (Gibco or Gemini Bio-Products) at 37°C in a humidified atmosphere with 5% CO2.

### Antibodies

The antibodies used in the study include: polyclonal rabbit anti-RBBP6 antibody (Origene Technologies, TA309830), polyclonal rabbit anti-hnRNPL (Abcam, ab32680), monoclonal rabbit anti-hnRNPUL1 (Abcam, ab134954), monoclonal rabbit anti-PEG10 (Abcam, ab215035), monoclonal mouse anti-FLAG M2 antibody (Sigma Aldrich, F1804), polyclonal rabbit anti-FLAG antibody (Sigma Aldrich, F7425), monoclonal mouse anti-HA antibody (Sigma Aldrich, H3663), polyclonal rabbit anti-HA antibody (Sigma Aldrich, H6908), monoclonal mouse anti-β-tubulin antibody (Sigma Aldrich, T8328), anti-Ebola GP antibody clone 4F3 (IBT Bioservices, 0201-020), goat IgG (H+L) anti-mouse Alexa Flour 546 conjugate (ThermoFisher scientific, A11030), rabbit anti-peptide serum raised against EBOV VP30 (2-25aa) and NP (97-119aa) and anti-VP30 pSer VP30 (Lier et al., 2017).

### Plasmid Constructs

The coding sequences of human hnRNPL (NM_001533.2), hnRNPUL1 (NM_007040) and PEG10 (NM_015068.3) were PCR amplified from ORF clones (hnRNPL, GenScript #OHu14072; hnRNPUL1, Origene #RC200576; PEG10, GenScript #OHu101111) and the products were cloned into the mammalian expression plasmid pCAGGS. The amino acid regions containing PPxPxY motifs in hnRNPL (peptide_1: 325-350aa; peptide_2: 360-385aa), hnRNPUL1 (peptide_1: 702-727aa; peptide_2: 742-767aa; peptide_3: 769-794aa) and PEG10 (peptide_1: 672-697aa; peptide_2: 690-708aa) were cloned as fusions to an N-terminal GFP in pCAGGS.

All expression plasmids for human RBBP6, GFP-RBBP6 peptide, EBOV minigenome assay and all VP30 mutant constructs (E197A, D202A, E197A/D202A/Q203A, Q229A, W230A, Q229A/W230A) have been previously described (Edwards, Pietzsch et al., 2015, Xu et al., 2017). The sequences of the constructs were validated by Sanger DNA sequencing prior to use.

### Co-immunoprecipitation Assays

HEK293T cells were transfected with the indicated plasmids using Lipofectamine 2000 (Invitrogen) as per the manufacturer's instructions. 24 h post-transfection, cells were harvested in NP-40 lysis buffer (50mM Tris-Cl pH 7.5, 280 mM NaCl, 0.5% NP-40, 2mM EDTA, 10% glycerol) supplemented with cOmplete mini protease inhibitor tablets (Roche). Clarified cell lysates were incubated with anti-GFP (MBL International Corp.), -HA or -FLAG magnetic beads (Sigma-Aldrich) for 1-2 h at 4°C, followed by washing the magnetic beads with NP-40 lysis buffer five times. Protein complexes were eluted by direct incubation in 1X SDS loading buffer and heating at 95°C. Eluates and whole-cell lysates were analyzed by western blotting using the indicated antibodies.

### Minigenome Assays

HEK293T cells were transfected with expression plasmids encoding EBOV proteins VP35, L, NP, VP30 along with T7 RNA polymerase and a plasmid that produces the negative-sense minigenome RNA encoding *Renilla* luciferase flanked by *cis*-acting transcription and replication signals from the EBOV genome. A plasmid expressing firefly luciferase from an RNA polymerase II promoter was included as an internal control. Forty-eight hours post-transfection, reporter activity was measured using the Dual-luciferase Reporter assay system (Promega). *Renilla* luciferase activity was normalized to firefly luciferase values and plotted as fold MG activity calculated relative to no VP30 control.

### Gene Silencing using siRNA

For minigenome assays, HEK293T cells were transfected with 25nM of SMARTpool:Accell siRNA targeting RBBP6 (Dharmacon), or 10nM siRNA duplexes targeting hnRNPL (QIAgen), hnRNPUL1 (ThermoFisher Scientific) or PEG10 (QIAgen) (final concentration made up to 25nM with control siRNA) or 25nM control ON-TARGET plus Non-targeting siRNA (ThermoFisher Scientific) using lipofectamine RNAiMAX (Invitrogen). Twenty-four hours post-siRNA transfection, cells were transfected with plasmids for the minigenome assay and luciferase activity was assessed as above.

### Hydrogen Deuterium Exchange

VP30 (130-272) was incubated individually with host factor peptides RBBP6, hnRNPUL1_1, hnRNPUL1_2, hnRNPUL1_3, hnRNPL_2 and PEG10_2 in the 10mM HEPES Buffer (150 mM NaCl, pH = 7.0) on ice for 45 mins. RBBP6 was at a 4:1 ratio to VP30 (130-272) and other host factor peptides were at 20:1 ratio. An aliquot of 2 μL of eVP30 stock solution (40 μM for bound or unbound) was added to 18 μL labeling buffer (10 mM HEPES buffer, 150 mM NaCl in 90% D2O, pH=7.0) and incubated on ice for 10 s, 30 s, 1 min, 2 mins, 15 mins, 2 h and 4 h. Quenching buffer of 30 μL (50 mM TCEP, 4 M urea in 10 mM HEPES, pH = 2.5) was then added to stop the deuterium exchange for 30 s on ice. Samples were then snap-frozen immediately with liquid nitrogen and kept at −80 °C until mass spectrometric analysis. All measurements were performed in triplicate. Before mass spectrometry, each sample vial was incubated at 25 °C for 30 s to thaw, followed by injection into a custom-built HDX platform. Through a ZORBAX Eclipse XDB C8 column (2.1 mm × 15 mm, Agilent Technologies), protein samples were desalted with 0.1% trifluoroacetic acid in H2O for 3 mins. On-line digestion was then conducted on an immobilized pepsin column (2 mm × 20 mm) at 200 μL/min flow rate. A Hypersil Gold C18 column (2.1 mm × 50 mm, ThermoFisher Scientific) with a 9.5-min-linear gradient (4–40% acetonitrile with 0.1% formic acid, 200 μL/min flow rate) was used for peptide separation right followed by LTQ-FT mass spectrometry analysis (ThermoFisher Scientific) with data acquisition at a mass resolving power of 100,000 at m/z 400. Residue-level HDX data were analyzed and exported from HDExaminer (Sierra Analytics, v2.5.0) by using “heavy” heat map smoothing to show HDX at the peptide and residue levels.

### RNA Isolation and Quantitative RT-PCR

HEK293T cells were transfected with the components of the EBOV minigenome system. Cells were harvested 48 h post-transfection and total RNA was isolated from the cells using the RNeasy mini kit (QIAgen) as per the manufacturer’s protocol. Residual plasmid contamination was removed by treatment with ezDNase enzyme (Invitrogen). For cDNA synthesis, ~1μg of RNA was used for reverse transcription of either positive strand (cRNA/mRNA) or negative strand RNA (vRNA) using SuperScript® IV First-Strand Synthesis kit (ThermoFisher Scientific). Subsequently, real-time PCR was performed using PerfeCTa SYBR Green FastMix (QuantaBio) on a CFX96 Real-Time PCR detection system (Bio-Rad), with PCR conditions as follows: initial denaturation at 95°C for 30s and then 40 cycles of 95°C for 5s, 55°C for 15s and 68°C for 10s. Expression levels were normalized to the β-actin control. Percent transcription or replication was calculated and set relative to the empty vector control sample.

### EBOV Infection Assays

All studies with infectious EBOV were performed under biosafety level 4 containment at Texas Biomedical Research Institute or the National Emerging Infectious Diseases Laboratory (NEIDL) at Boston University. For siRNA knockdown studies, 2×10^4^ siRNA-treated HeLa cells were plated into 96-well plates in triplicate for infection with EBOV-EGFP. Cells were allowed to adhere for 24 hours, followed by virus infection at an MOI of either 0.05 or 0.5 for 24 h. Cells were fixed in 10% buffered formalin and stained with the Hoechst dye. Relative infection rate was calculated as the ratio of EGFP-positive cells to cell nuclei by analyzing ≥7,000 cells, in triplicates. siRNA-mediated knockdown was assessed by western blotting in the cell lysates collected at the same time as infection. Three independent experiments were performed for each siRNA tested.

### Isolation of proteins from EBOV-infected lysates

HeLa cells were infected with EBOV-GFP at an MOI of 0.1. Twenty-four hours post-infection, cells were lysed in TRIzol reagent and proteins were isolated as described (Batra et al., 2018).

### Statistical analysis

All statistical analysis was performed using GraphPad Prism 7, using appropriate parameters as indicated in the figure legends and text. Data values were considered significantly different if the p-value was <0.05. Unless stated otherwise, ANOVA with Tukey’s multiple comparisons test was used.

## ACKNOWLEDGEMENTS

This study was supported by NIH grants R01AI143292 to GKA, CFB and NJK and P01AI120943 to GKA, CFB, DWL, MLG, and RAD. It was also supported by Department of the Defense, Defense Threat Reduction Agency grant HDTRA1-16-1-0033 to C.F.B. and G.K.A.. The mass spectrometry was supported by NIGMS of the NIH (Grant P41GM103422). We thank Megan R. Edwards for critical reading of the manuscript.

## AUTHOR CONTRIBUTIONS

J.B., O.S., G.I.S, M.A., M.Z., D.L. and C.G.W conducted the experiments; J.B., O.S., G.I.S., M.A., M.Z., D.L., M.L.G., R.A.D., D.W.L., G.K.A., N.J.K. and C.F.B. analyzed the data; J.B., O.S., M.A., M.L.G., D.W.L., R.A.D., G.K.A., N.J.K. and C.F.B. designed the experiments; N.B. and S.B. provided critical reagents; J.B., G.K.A. and C.F.B. wrote the paper with input from all co-authors.

## EXPANDED VIEW FIGURE LEGENDS

**Figure EV1. Co-IP between VP30 and endogenous PEG10.** Huh7 cells were transfected with empty vector (-lane) or FLAG-VP30 expression plasmid (+ lane). IPs (IP: Anti-FLAG) and whole cell lysates (WCL) were analyzed by immunoblotting with anti-FLAG or anti-PEG10 antibodies.

**Figure EV2. Effects of PEG10 knockdown on EBOV RNA synthesis. (A)** Minigenome activity upon knockdown of endogenous PEG10. Huh7 cells were transfected with scrambled siRNA or siRNA targeting PEG10. Twenty-four hours post-transfection, cells were transfected with MG assay plasmids. Data represent mean ± S.D. from one representative experiment (n=3) of at least two independent experiments. **(B)** Immunoblots detecting the proteins levels of PEG10 and VP30 are shown.

**Figure EV3. Binding to VP30 of PPxPxY motifs from different proteins. (A)** Table summarizing VP30-peptide interactions and activity, based on co-IP, FP assay and inhibition of MG activity. Percent inhibition +++: >80%; ++: 60-80%; +: 40-60%. **(B)** Co-immunoprecipitation assay to assess HA-VP40 interaction with FLAG-VP30. HEK293T cells were transfected with HA-VP40 plasmid and either empty vector or FLAG-VP30 plasmid. An anti-FLAG IP was performed. IP and WCL were analyzed by western blotting with anti-HA and anti-FLAG antibodies with anti-β-tubulin providing a loading control. **(C)** Anti-GFP immunoprecipitation of HA-VP30 with wild-type or mutated VP40 peptides fused to GFP. GFP without a fusion partner served as a control. Immunoblots with anti-HA, anti-GFP and anti-β-tubulin are shown. **(D)** GFP fused to peptides derived from human and bat (*Rousettus aegyptiacus*) RBBP6 were expressed along with HA-VP30. Co-IP was performed using anti-GFP magnetic beads and representative immunoblots for IP and WCL are shown. **(E)** Equilibrium dissociation curves of FITC-RBBP6 peptide to eVP30 (130-272) as it is outcompeted by increasing concentrations (0.13-500 μM) of human and bat RBBP6 peptides. Fluorescence polarization was determined with constant concentrations of FITC-RBBP6 and eVP30 (130-272), at 0.50 μM and 3.8 μM respectively. Experiments were performed in two independent duplicates. Error bars represent standard deviation. **(F)** Minigenome activity upon titration of GFP fused to human and bat-derived RBBP6 peptides. GFP alone was used as a control. Reporter activity was read at 48h post-transfection and fold MG activity was calculated relative to a no VP30 control. Data represent the mean ± S.D. from one representative experiment (n=3) of at least two independent experiments. Statistical significance was calculated relative to GFP control for each concentration tested. ****p<0.00005; ***p<0.0005, **p<0.005, *p<0.05

**Figure EV4. Detailed HDX interaction data. (A)** HDX kinetic plots for all VP30 peptides. **(B)** Statistical analysis of deuterium uptake differences of all time points confirms the binding regions. Deuterium uptake differences between different bound VP30 and unbound VP30 were calculated for each time point from 10 s to 4 h, depicted by gradient colors. Standard deviation between triplicates was calculated separately for each time point representing the bound and unbound states. Plotted on the graph is a 3-fold propagation error of each time point as shown by the error bars, giving 99.7% certainty on the observed difference. VP30 bound with RBBP6, hnRNPUL1_1, hnRNPUL1_3, hnRNPL_2 and PEG10_2 showed the same binding regions and similar regions exhibiting remote conformational or dynamics reduction owing to binding. There is no significant difference for VP30 with hnRNPUL1_2; therefore, it served as a negative control.

**Figure EV5. VP30 mutants interaction with hnRNPUL1 and PEG10. (A, B)** Representative western blots of co-IP experiments are presented to assess interaction between the indicated VP30 mutants and hnRNPUL1 (A) or PEG10 (B).

---

**Figure S1. Western blots assessing levels of hnRNP L and hnRNPUL1 in knockdown experiments.** HeLa cells were mock-transfected (Mock), transfected with 5 nM of scrambled siRNA (SCR si) and siRNAs targeting hnRNP L (A) or hnRNPUL1 (B). Seventy-two hours post-transfection, western blots were performed on whole cell lysates using antibodies to the indicated proteins.

## REFERENCES

1. Ammosova T, Pietzsch CA, Saygideger Y, Ilatovsky A, Lin X, Ivanov A, Kumari N, Jerebtsova M, Kulkarni A, Petukhov M, Uren A, Bukreyev A, Nekhai S (2018) Protein Phosphatase 1-Targeting Small-Molecule C31 Inhibits Ebola Virus Replication. J Infect Dis 218: S627–S635

2. Ball LJ, Kuhne R, Schneider-Mergener J, Oschkinat H (2005) Recognition of proline-rich motifs by protein-protein-interaction domains. Angew Chem Int Ed Engl 44: 2852–69

3. Batra J, Hultquist JF, Liu D, Shtanko O, Von Dollen J, Satkamp L, Jang GM, Luthra P, Schwarz TM, Small GI, Arnett E, Anantpadma M, Reyes A, Leung DW, Kaake R, Haas P, Schmidt CB, Schlesinger LS, LaCount DJ, Davey RA et al. (2018) Protein Interaction Mapping Identifies RBBP6 as a Negative Regulator of Ebola Virus Replication. Cell 175: 1917–1930 e13

4. Bausch DG, Rojek A (2016) West Africa 2013: Re-examining Ebola. Microbiol Spectr 4

5. Becker S, Rinne C, Hofsass U, Klenk HD, Muhlberger E (1998) Interactions of Marburg virus nucleocapsid proteins. Virology 249: 406–17

6. Biedenkopf N, Hartlieb B, Hoenen T, Becker S (2013) Phosphorylation of Ebola virus VP30 influences the composition of the viral nucleocapsid complex: impact on viral transcription and replication. J Biol Chem 288: 11165–74

7. Biedenkopf N, Lier C, Becker S (2016a) Dynamic Phosphorylation of VP30 Is Essential for Ebola Virus Life Cycle. J Virol 90: 4914–4925

8. Biedenkopf N, Schlereth J, Grunweller A, Becker S, Hartmann RK (2016b) RNA Binding of Ebola Virus VP30 Is Essential for Activating Viral Transcription. J Virol 90: 7481–96

9. Cao L, Liu S, Li Y, Yang G, Luo Y, Li S, Du H, Zhao Y, Wang D, Chen J, Zhang Z, Li M, Ouyang S, Gao X, Sun Y, Wang Z, Yang L, Lin R, Wang P, You F (2019) The Nuclear Matrix Protein SAFA Surveils Viral RNA and Facilitates Immunity by Activating Antiviral Enhancers and Super-enhancers. Cell Host Microbe 26: 369–384 e8

10. Carette JE, Raaben M, Wong AC, Herbert AS, Obernosterer G, Mulherkar N, Kuehne AI, Kranzusch PJ, Griffin AM, Ruthel G, Dal Cin P, Dye JM, Whelan SP, Chandran K, Brummelkamp TR (2011) Ebola virus entry requires the cholesterol transporter Niemann-Pick C1. Nature 477: 340–3

11. Edwards MR, Pietzsch C, Vausselin T, Shaw ML, Bukreyev A, Basler CF (2015) High-Throughput Minigenome System for Identifying Small-Molecule Inhibitors of Ebola Virus Replication. ACS Infect Dis 1: 380–7

12. Enterlein S, Volchkov V, Weik M, Kolesnikova L, Volchkova V, Klenk HD, Muhlberger E (2006) Rescue of recombinant Marburg virus from cDNA is dependent on nucleocapsid protein VP30. J Virol 80: 1038–43

13. Gurunathan G, Yu Z, Coulombe Y, Masson JY, Richard S (2015) Arginine methylation of hnRNPUL1 regulates interaction with NBS1 and recruitment to sites of DNA damage. Sci Rep 5: 10475

14. Halfmann P, Kim JH, Ebihara H, Noda T, Neumann G, Feldmann H, Kawaoka Y (2008) Generation of biologically contained Ebola viruses. Proc Natl Acad Sci U S A 105: 1129–33

15. Han Z, Sagum CA, Bedford MT, Sidhu SS, Sudol M, Harty RN (2016) ITCH E3 Ubiquitin Ligase Interacts with Ebola Virus VP40 To Regulate Budding. J Virol 90: 9163–71

16. Han Z, Sagum CA, Takizawa F, Ruthel G, Berry CT, Kong J, Sunyer JO, Freedman BD, Bedford MT, Sidhu SS, Sudol M, Harty RN (2017) Ubiquitin Ligase WWP1 Interacts with Ebola Virus VP40 To Regulate Egress. J Virol 91

17. Harty RN, Brown ME, Wang G, Huibregtse J, Hayes FP (2000) A PPxY motif within the VP40 protein of Ebola virus interacts physically and functionally with a ubiquitin ligase: implications for filovirus budding. Proc Natl Acad Sci U S A 97: 13871–6

18. Hishida T, Naito K, Osada S, Nishizuka M, Imagawa M (2007) peg10, an imprinted gene, plays a crucial role in adipocyte differentiation. FEBS Lett 581: 4272–8

19. Hong Z, Jiang J, Ma J, Dai S, Xu T, Li H, Yasui A (2013) The role of hnRPUL1 involved in DNA damage response is related to PARP1. PLoS One 8: e60208

20. Ilinykh PA, Tigabu B, Ivanov A, Ammosova T, Obukhov Y, Garron T, Kumari N, Kovalskyy D, Platonov MO, Naumchik VS, Freiberg AN, Nekhai S, Bukreyev A (2014) Role of protein phosphatase 1 in dephosphorylation of Ebola virus VP30 protein and its targeting for the inhibition of viral transcription. J Biol Chem 289: 22723–38

21. Ilunga Kalenga O, Moeti M, Sparrow A, Nguyen VK, Lucey D, Ghebreyesus TA (2019) The Ongoing Ebola Epidemic in the Democratic Republic of Congo, 2018-2019. N Engl J Med 381: 373–383

22. John SP, Wang T, Steffen S, Longhi S, Schmaljohn CS, Jonsson CB (2007) Ebola virus VP30 is an RNA binding protein. J Virol 81: 8967–76

23. Kirchdoerfer RN, Moyer CL, Abelson DM, Saphire EO (2016) The Ebola Virus VP30-NP Interaction Is a Regulator of Viral RNA Synthesis. PLoS Pathog 12: e1005937

24. Kruse T, Biedenkopf N, Hertz EPT, Dietzel E, Stalmann G, Lopez-Mendez B, Davey NE, Nilsson J, Becker S (2018) The Ebola Virus Nucleoprotein Recruits the Host PP2A-B56 Phosphatase to Activate Transcriptional Support Activity of VP30. Mol Cell 69: 136–145 e6

25. Liang J, Sagum CA, Bedford MT, Sidhu SS, Sudol M, Han Z, Harty RN (2017) Chaperone-Mediated Autophagy Protein BAG3 Negatively Regulates Ebola and Marburg VP40-Mediated Egress. PLoS Pathog 13: e1006132

26. Licata JM, Simpson-Holley M, Wright NT, Han Z, Paragas J, Harty RN (2003) Overlapping motifs (PTAP and PPEY) within the Ebola virus VP40 protein function independently as late budding domains: involvement of host proteins TSG101 and VPS-4. J Virol 77: 1812–9

27. Lier C, Becker S, Biedenkopf N (2017) Dynamic phosphorylation of Ebola virus VP30 in NP-induced inclusion bodies. Virology 512: 39–47

28. Martin-Serrano J, Eastman SW, Chung W, Bieniasz PD (2005) HECT ubiquitin ligases link viral and cellular PPXY motifs to the vacuolar protein-sorting pathway. J Cell Biol 168: 89–101

29. Martinez MJ, Biedenkopf N, Volchkova V, Hartlieb B, Alazard-Dany N, Reynard O, Becker S, Volchkov V (2008) Role of Ebola virus VP30 in transcription reinitiation. J Virol 82: 12569–73

30. Martinez MJ, Volchkova VA, Raoul H, Alazard-Dany N, Reynard O, Volchkov VE (2011) Role of VP30 phosphorylation in the Ebola virus replication cycle. J Infect Dis 204 Suppl 3: S934–40

31. Mehedi M, Hoenen T, Robertson S, Ricklefs S, Dolan MA, Taylor T, Falzarano D, Ebihara H, Porcella SF, Feldmann H (2013) Ebola virus RNA editing depends on the primary editing site sequence and an upstream secondary structure. PLoS Pathog 9: e1003677

32. Modrof J, Becker S, Muhlberger E (2003) Ebola virus transcription activator VP30 is a zinc-binding protein. J Virol 77: 3334–8

33. Modrof J, Muhlberger E, Klenk HD, Becker S (2002) Phosphorylation of VP30 impairs ebola virus transcription. J Biol Chem 277: 33099–104

34. Muhlberger E, Lotfering B, Klenk HD, Becker S (1998) Three of the four nucleocapsid proteins of Marburg virus, NP, VP35, and L, are sufficient to mediate replication and transcription of Marburg virus-specific monocistronic minigenomes. J Virol 72: 8756–64

35. Muhlberger E, Weik M, Volchkov VE, Klenk HD, Becker S (1999) Comparison of the transcription and replication strategies of marburg virus and Ebola virus by using artificial replication systems. J Virol 73: 2333–42

36. Nanbo A, Watanabe S, Halfmann P, Kawaoka Y (2013) The spatio-temporal distribution dynamics of Ebola virus proteins and RNA in infected cells. Sci Rep 3: 1206

37. Nsio J, Kapetshi J, Makiala S, Raymond F, Tshapenda G, Boucher N, Corbeil J, Okitandjate A, Mbuyi G, Kiyele M, Mondonge V, Kikoo MJ, Van Herp M, Barboza P, Petrucci R, Benedetti G, Formenty P, Muyembe Muzinga B, Ilunga Kalenga O, Ahuka S et al. (2019) 2017 Outbreak of Ebola Virus Disease in Northern Democratic Republic of Congo. J Infect Dis

38. Ntwasa M (2016) Retinoblastoma Binding Protein 6, Another p53 Monitor. Trends Cancer 2: 635–637

39. Peng YP, Zhu Y, Yin LD, Zhang JJ, Wei JS, Liu X, Liu XC, Gao WT, Jiang KR, Miao Y (2017) PEG10 overexpression induced by E2F-1 promotes cell proliferation, migration, and invasion in pancreatic cancer. J Exp Clin Cancer Res 36: 30

40. Schlereth J, Grunweller A, Biedenkopf N, Becker S, Hartmann RK (2016) RNA binding specificity of Ebola virus transcription factor VP30. RNA Biol 13: 783–98

41. Seo JY, Kim DY, Kim SH, Kim HJ, Ryu HG, Lee J, Lee KH, Kim KT (2017) Heterogeneous nuclear ribonucleoprotein (hnRNP) L promotes DNA damage-induced cell apoptosis by enhancing the translation of p53. Oncotarget 8: 51108–51122

42. Takamatsu Y, Krahling V, Kolesnikova L, Halwe S, Lier C, Baumeister S, Noda T, Biedenkopf N, Becker S (2020) Serine-Arginine Protein Kinase 1 Regulates Ebola Virus Transcription. mBio 11

43. Tigabu B, Ramanathan P, Ivanov A, Lin X, Ilinykh PA, Parry CS, Freiberg AN, Nekhai S, Bukreyev A (2018) Phosphorylated VP30 of Marburg Virus Is a Repressor of Transcription. J Virol 92

44. Timmins J, Schoehn G, Ricard-Blum S, Scianimanico S, Vernet T, Ruigrok RW, Weissenhorn W (2003) Ebola virus matrix protein VP40 interaction with human cellular factors Tsg101 and Nedd4. J Mol Biol 326: 493–502

45. Weik M, Modrof J, Klenk HD, Becker S, Muhlberger E (2002) Ebola virus VP30-mediated transcription is regulated by RNA secondary structure formation. J Virol 76: 8532–9

46. Xie T, Pan S, Zheng H, Luo Z, Tembo KM, Jamal M, Yu Z, Yu Y, Xia J, Yin Q, Wang M, Yuan W, Zhang Q, Xiong J (2018) PEG10 as an oncogene: expression regulatory mechanisms and role in tumor progression. Cancer Cell Int 18: 112

47. Xu W, Luthra P, Wu C, Batra J, Leung DW, Basler CF, Amarasinghe GK (2017) Ebola virus VP30 and nucleoprotein interactions modulate viral RNA synthesis. Nat Commun 8: 15576

48. Yasuda J, Nakao M, Kawaoka Y, Shida H (2003) Nedd4 regulates egress of Ebola virus-like particles from host cells. J Virol 77: 9987–92

49. Zhou X, Li Q, He J, Zhong L, Shu F, Xing R, Lv D, Lei B, Wan B, Yang Y, Wu H, Mao X, Zou Y (2017) HnRNP-L promotes prostate cancer progression by enhancing cell cycling and inhibiting apoptosis. Oncotarget 8: 19342–19353

