## Supplemental figures for "Non-canonical proline-tyrosine interactions with multiple host proteins regulate Ebola virus infection"

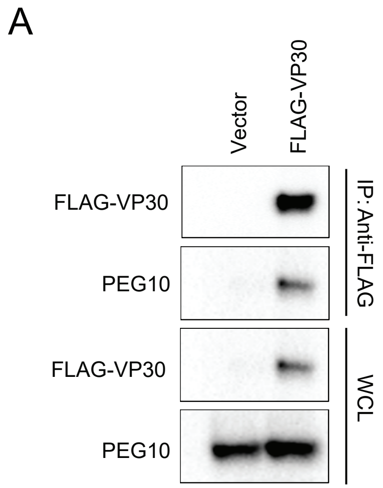


Figure EV1. Co-IP between VP30 and endogenous PEG10. Huh7 cells were transfected with empty vector (- lane) or FLAG-VP30 expression plasmid (+ lane). IPs (IP: Anti-FLAG) and whole cell lysates (WCL) were analyzed by immunoblotting with anti-FLAG or anti-PEG10 antibodies.

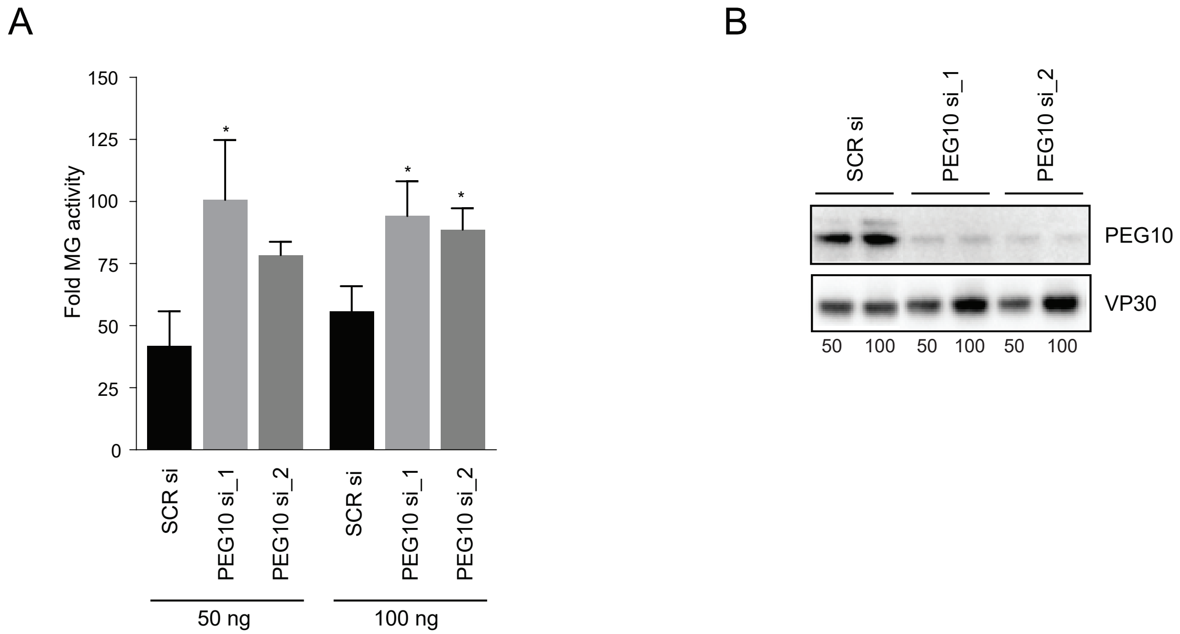


Figure EV2. Effects of PEG10 knockdown on EBOV RNA synthesis. (A) Minigenome activity upon knockdown of endogenous PEG10. Huh7 cells were transfected with scrambled siRNA or siRNA targeting PEG10. Twenty-four hours post-transfection, cells were transfected with MG assay plasmids. Data represent mean ± S.D. from one representative experiment (n=3) of at least two independent experiments. (B) Immunoblots detecting the proteins levels of PEG10 and VP30 are shown.

A

| Peptides | Co-IP | FPA | ↓MG |
|---|---|---|---|
| RBBP6 | +++ | +++ | +++ |
| hnRNP L_1 | - | - | - |
| hnRNP L_2 | + | + | + |
| hnRNPUL1_1 | +/- | + | - |
| hnRNPUL1_2 | - | - | - |
| hnRNPUL1_3 | ++ | + | ++ |
| PEG10_1 | - | - | + |
| PEG10_2 | ++ | + | +/- |

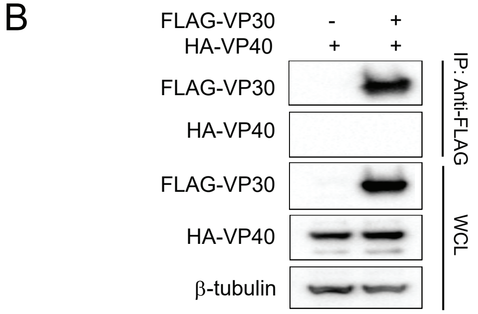


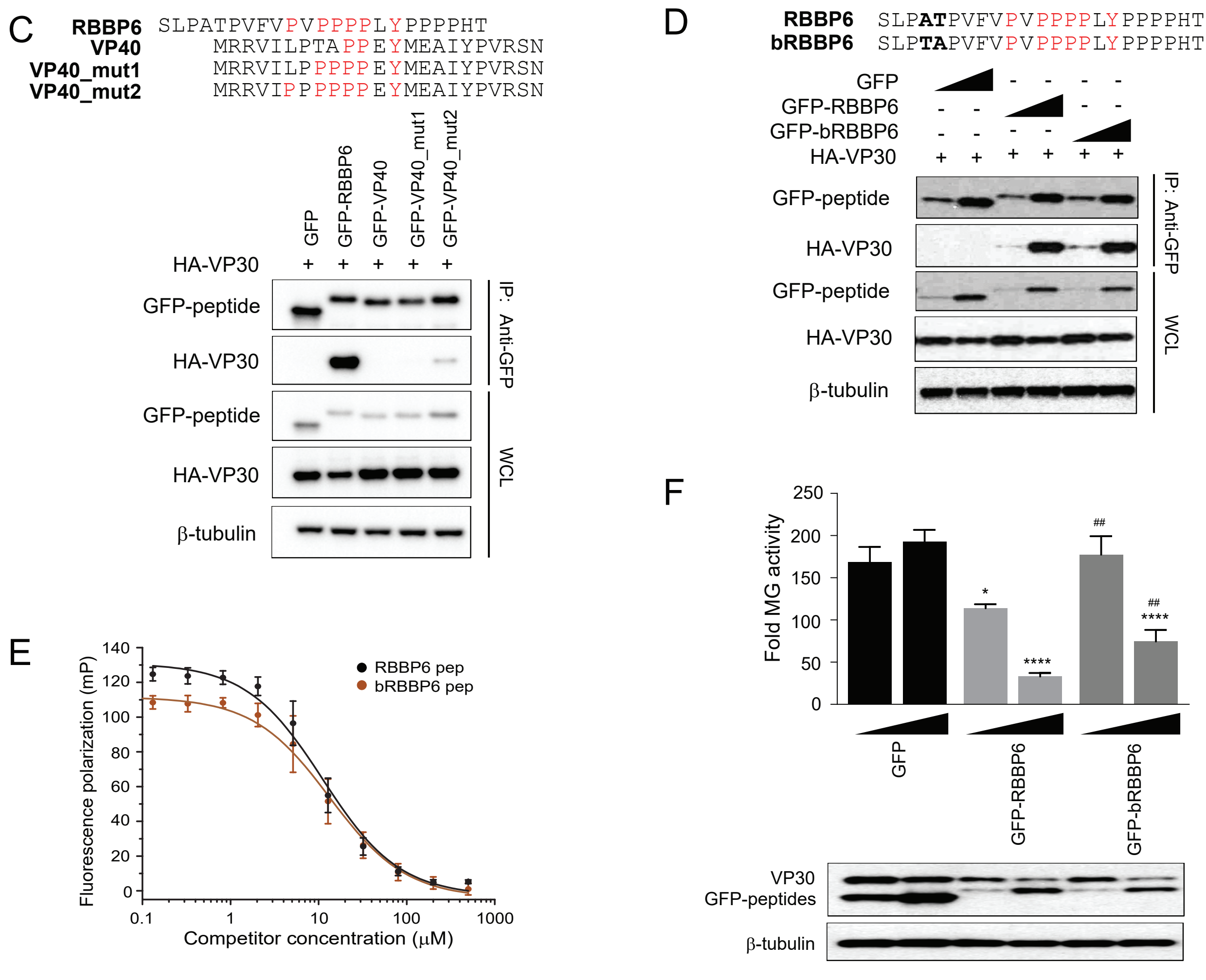


Figure EV3. Binding to VP30 of PPxPxY motifs from different proteins. (A) Table summarizing VP30-peptide interactions and activity, based on co-IP, FP assay and inhibition of MG activity. Percent inhibition +++: >80%; ++: 60-80%; +: 40-60%. (B) Co-immunoprecipitation assay to assess HA-VP40 interaction with FLAG-VP30. HEK293T cells were transfected with HA-VP40 plasmid and either empty vector or FLAG-VP30 plasmid. An anti-FLAG IP was performed. IP and WCL were analyzed by western blotting with anti-HA and anti-FLAG antibodies with anti-β-tubulin providing a loading control. (C) Anti-GFP immunoprecipitation of HA-VP30 with wild-type or mutated VP40 peptides fused to GFP. GFP without a fusion partner served as a control. Immunoblots with anti-HA, anti-GFP and anti-β-tubulin are shown. (D) GFP fused to peptides derived from human and bat (Rousettus aegyptiacus) RBBP6 were expressed along with HA-VP30. Co-IP was performed using anti-GFP magnetic beads and representative immunoblots for IP and WCL are shown. (E) Equilibrium dissociation curves of FITC-RBBP6 peptide to eVP30 (130-272) as it is outcompeted by increasing concentrations (0.13-500 μM) of human and bat RBBP6 peptides. Fluorescence polarization was determined with constant concentrations of FITC-RBBP6 and eVP30 (130-272), at 0.50 μM and 3.8 μM respectively. Experiments were performed in two independent duplicates. Error bars represent standard deviation. (F) Minigenome activity upon titration of GFP fused to human and bat-derived RBBP6 peptides. GFP alone was used as a control. Reporter activity was read at 48h post-transfection and fold MG activity was calculated relative to a no VP30 control. Data represent the mean ± S.D. from one representative experiment (n=3) of at least two independent experiments. Statistical significance was calculated relative to GFP control for each concentration tested. ****p<0.00005; ***p<0.0005, **p<0.005, *p<0.05

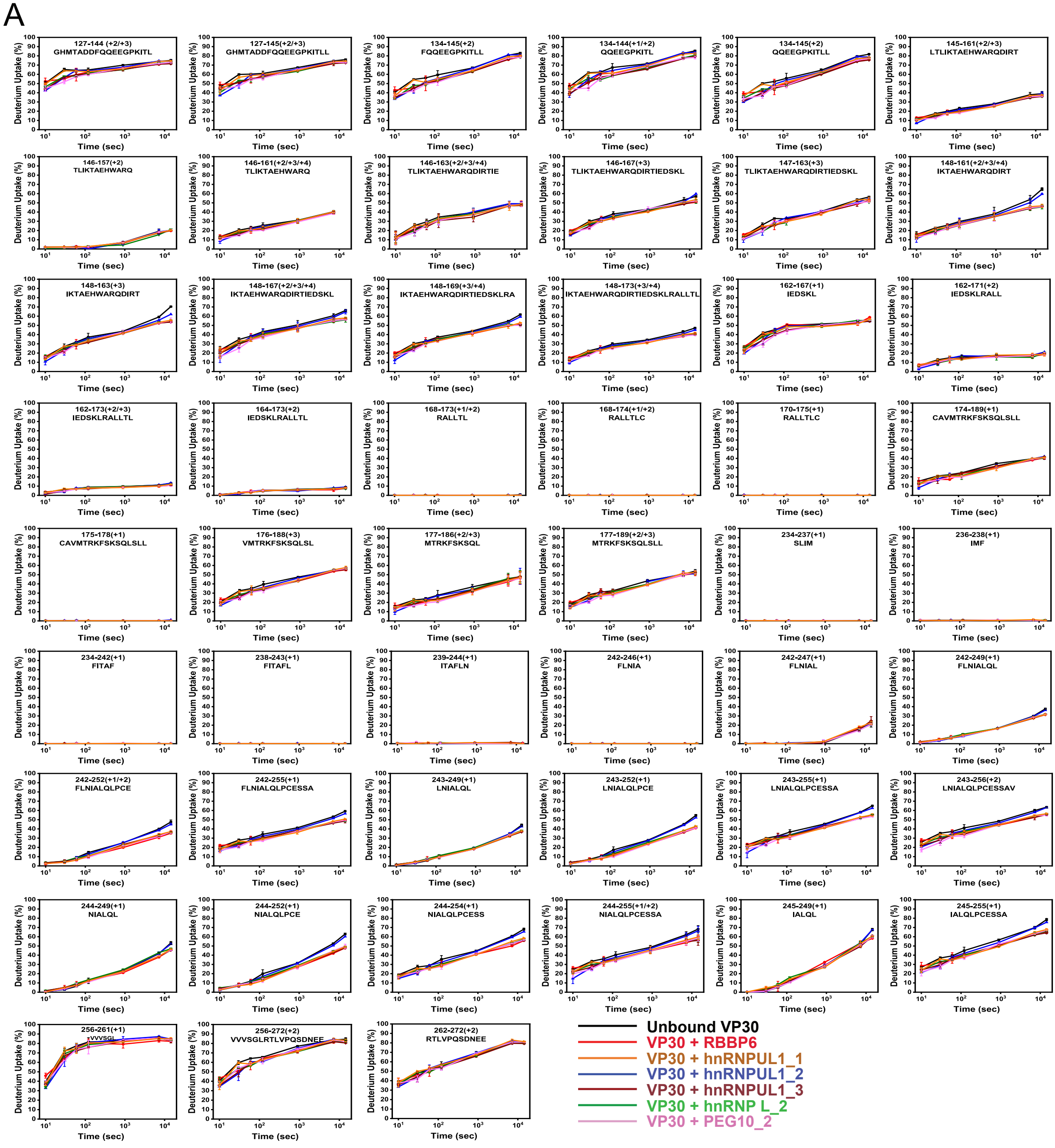
A
127-144 (+2/+3) GHMTADDFQQEEGPKITL
127-145(+2/+3) GHMTADDFQQEEGPKITLL
134-145(+2) FQQEEGPKITLL
134-144(+1/+2) QQEEGPKITL
134-145(+2) QQEEGPKITLL
145-161(+2/+3) LTLIKTAEHWARQDIRT
146-157(+2) TLIKTAEHWARQ
146-161(+2/+3/+4) TLIKTAEHWARQ
146-163(+2/+3/+4) TLIKTAEHWARQDIRTIE
146-167(+3) TLIKTAEHWARQDIRTIEDSKL
147-163(+3) TLIKTAEHWARQDIRTIEDSKL
148-161(+2/+3/+4) IKTAEHWARQDIRT
148-163(+3) IKTAEHWARQDIRT
148-167(+2/+3/+4) IKTAEHWARQDIRTIEDSKL
148-169(+3/+4) IKTAEHWARQDIRTIEDSKLRA
148-173(+3/+4) IKTAEHWARQDIRTIEDSKLRALLTL
162-167(+1) IEDSKL
162-171(+2) IEDSKLRALL
162-173(+2/+3) IEDSKLRALLTL
164-173(+2) IEDSKLRALLTL
168-173(+1/+2) RALLTL
168-174(+1/+2) RALLTLC
170-175(+1) RALLTLC
174-189(+1) CAVMTRKFSKSQLSLL
175-178(+1) CAVMTRKFSKSQLSLL
176-188(+3) VMTRKFSKSQLSL
177-186(+2/+3) MTRKFSKSQL
177-189(+2/+3) MTRKFSKSQLSLL
234-237(+1) SLIM
236-238(+1) IMF
234-242(+1) FITAF
238-243(+1) FITAFL
239-244(+1) ITAFLN
242-246(+1) FLNIA
242-247(+1) FLNIAL
242-249(+1) FLNIALQL
242-252(+1/+2) FLNIALQLPCE
242-255(+1) FLNIALQLPCESSA
243-249(+1) LNIALQL
243-252(+1) LNIALQLPCE
243-255(+1) LNIALQLPCESSA
243-256(+2) LNIALQLPCESSAV
244-249(+1) NIALQL
244-252(+1) NIALQLPCE
244-254(+1) NIALQLPCESS
244-255(+1/+2) NIALQLPCESSA
245-249(+1) IALQL
245-255(+1) IALQLPCESSA
256-261(+1) VVVSGL
256-272(+2) VVVSGLRTLVPQSDNEE
262-272(+2) RTLVPQSDNEE
Deuterium Uptake (%)
Time (sec)
Unbound VP30
VP30 + RBBP6
VP30 + hnRNPUL1_1
VP30 + hnRNPUL1_2
VP30 + hnRNPUL1_3
VP30 + hnRNP L_2
VP30 + PEG10_2

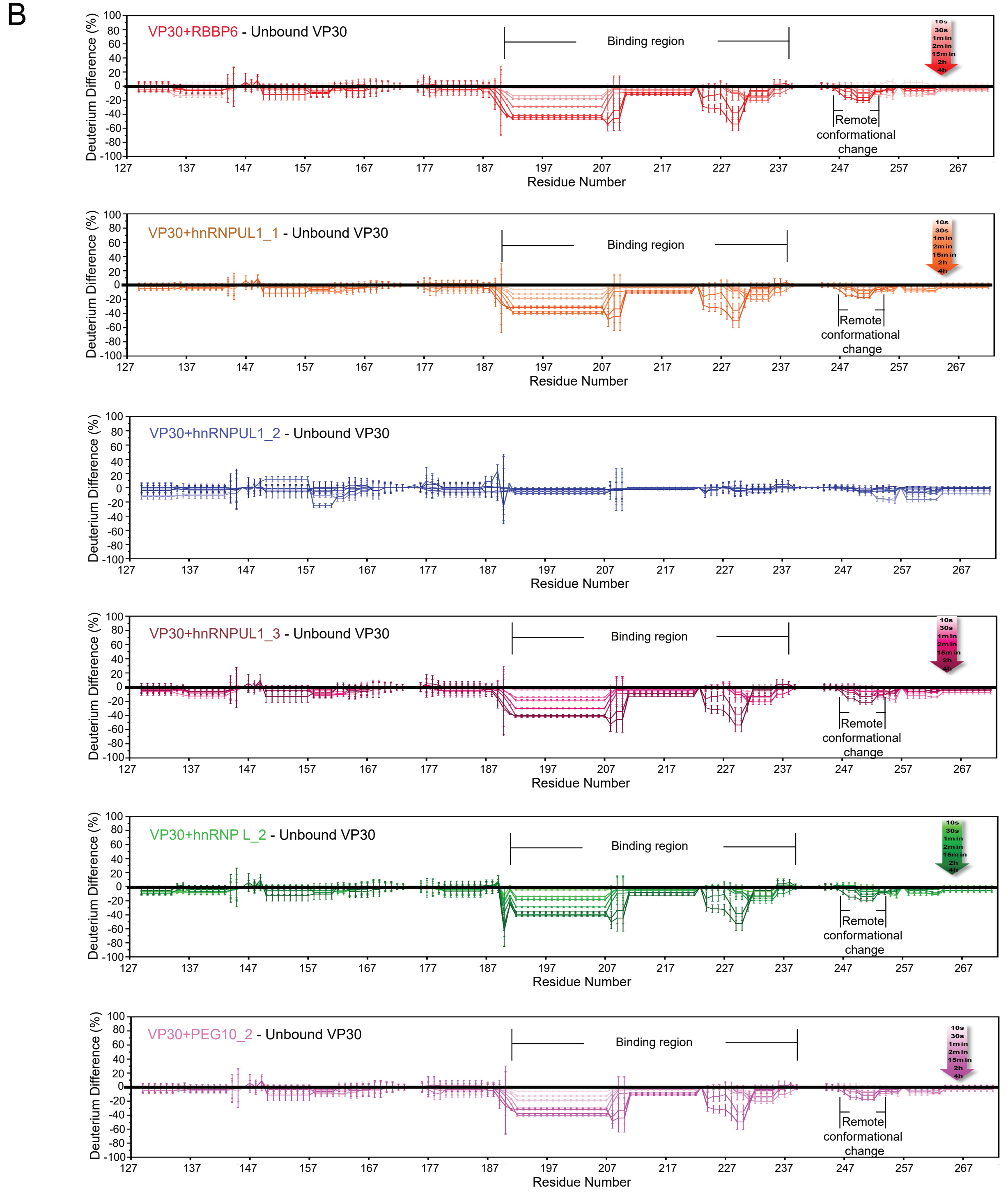


Figure EV4. Detailed HDX interaction data. (A) HDX kinetic plots for all VP30 peptides. (B) Statistical analysis of deuterium uptake differences of all time points confirms the binding regions. Deuterium uptake differences between different bound VP30 and unbound VP30 were calculated for each time point from 10 s to 4 h, depicted by gradient colors. Standard deviation between triplicates was calculated separately for each time point representing the bound and unbound states. Plotted on the graph is a 3-fold propagation error of each time point as shown by the error bars, giving 99.7% certainty on the observed difference. VP30 bound with RBBP6, hnRNPUL1_1, hnRNPUL1_3, hnRNPL_2 and PEG10_2 showed the same binding regions and similar regions exhibiting remote conformational or dynamics reduction owing to binding. There is no significant difference for VP30 with hnRNPUL1_2; therefore, it served as a negative control.

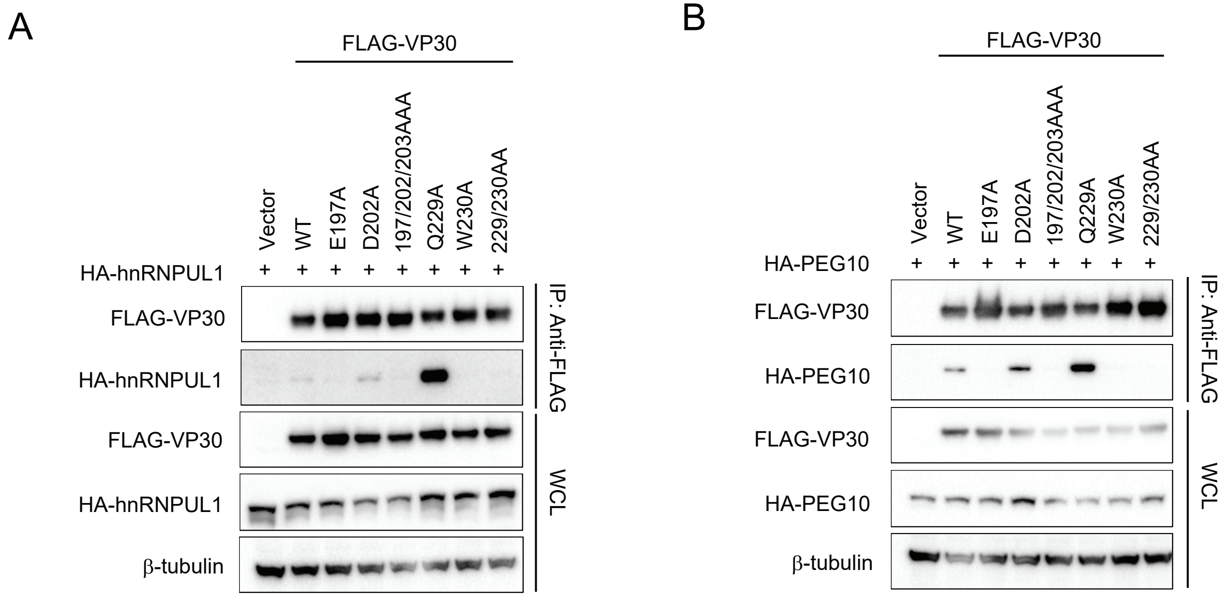


Figure EV5. VP30 mutants interaction with hnRNPUL1 and PEG10. (A, B) Representative western blots of co-IP experiments are presented to assess interaction between the indicated VP30 mutants and hnRNPUL1 (A) or PEG10 (B).

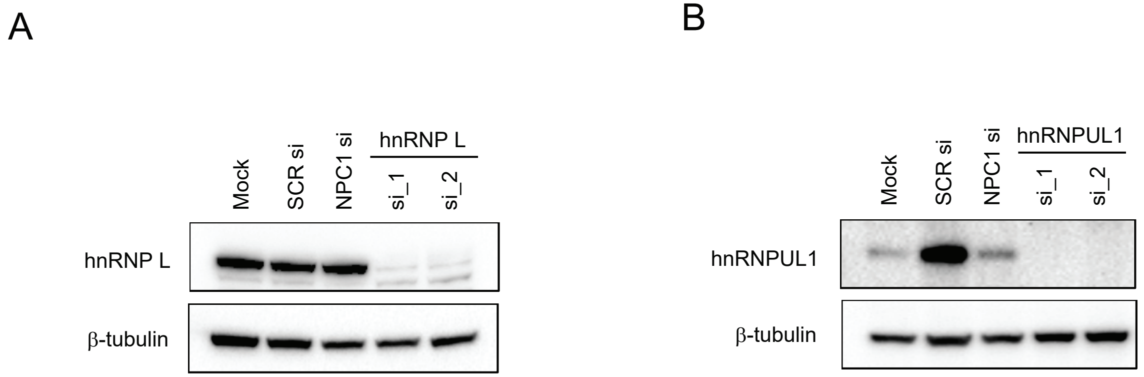


Appendix Figure S1. Western blots assessing levels of hnRNP L and hnRNPUL1 in knockdown experiments. HeLa cells were mock-transfected (Mock), transfected with 5 nM of scrambled siRNA (SCR si) and siRNAs targeting hnRNP L (A) or hnRNPUL1 (B). Seventy-two hours post-transfection, western blots were performed on whole cell lysates using antibodies to the indicated proteins.
